# Higher-Order Interactions in Neuronal Function: From Genes to Ionic Currents in Biophysical Models

**DOI:** 10.1101/2024.12.16.628700

**Authors:** Maria Reva, Alexis Arnaudon, Michael Zbili, Abdullah Makkeh, Henry Markram, Jean-Marc Goaillard, Werner Van Geit

**Author notes:** These authors share senior authorship (Jean-Marc Goaillard, Werner Van Geit).

## Abstract

Neuronal firing patterns are the consequence of precise variations in neuronal membrane potential, which are themselves shaped by multiple ionic currents. In this study, we use biophysical models, statistical methods, and information theory to explore the interaction between these ionic currents and neuron electrophysiological phenotype. We created numerous electrical models with diverse firing patterns. By analyzing these models, we identified intricate relationships between model parameters and electrical features. Our findings show that neuronal activity is often influenced by multiple biophysical model parameters, in a non-additive (i.e. synergistic) fashion. When comparing this with single-cell RNAseq data, we found a contrasting structure: gene expression profiles were dominated by redundancy, reflecting differences in regulatory constraints and sampling diversity. This research sheds light on the complex links between biophysical parameters and neuronal phenotypes.

---

While neurons are often categorized under broad classes, a number of studies have demonstrated that, even in seemingly homogeneous neuronal populations, neurons exhibit profound heterogeneity across multiple dimensions, ranging from genetic and molecular differences to variations in electrophysiological properties ^1,2,3,4,5^. Thus neuronal diversity appears as an intricate and multifaceted phenomenon, involving not only variations between neuronal types but also within neuronal types.

Genetic heterogeneity, for instance, is reflected in the differential expression of genes, particularly those involved in the regulation of ion channels. Interestingly, the landscape of ion channel genetic expression does not seem to be random. Several studies found correlations between the levels of expression of various ion channels ^6,7,8,9^, the patterns of these correlations giving rise to co-expression modules. Moreover, these coexpression modules appear to be specific of neuronal types. This suggests that, despite the variability in ion channel expression, neuronal phenotype might be defined by a specific co-expression module rather than by individual levels of expression of ion channels. A study utilizing single-cell multiplex RT-PCR on neocortical neurons identified 26 ion channel genes clustering with three calcium-binding proteins—calbindin, parvalbumin, and calretinin—suggesting a potential link between gene expression profiles and calcium dynamics ^10^. In fact, single-cell transcriptomics has consistently uncovered significant variations in mRNA and non-coding RNA levels between individual neurons of the same neuronal type, emphasizing the significance of molecular heterogeneity. Electrophysiological diversity adds yet another layer of complexity to neuronal phenotype. The combination of ion channel gene expression, along with the factors governing mRNA transcription, protein translation, and post-translational modifications, plays a crucial role in shaping the electrical phenotype of a neuron ^11^, such that variations in electrophysiological behaviors across neurons are often associated with genetic and molecular variations ^12,13,14,15^.

To decipher this complex link, tools like information theory offer valuable insights by helping researchers analyze large-scale data and distinguish meaningful heterogeneity from noise. For example, the Partial Information Decomposition for Cell regulatory networks (PIDC) enables the identification of gene regulatory networks (GRNs) from single-cell data, moving beyond pairwise correlations to uncover higher-order relationships ^16^. High-order interactions in complex systems such as gene regulatory networks can exhibit both synergistic and redundant characteristics. Synergistic interactions occur when the combined effect of multiple components is greater than the sum of their individual effects. This means that certain information is only accessible when considering the joint state of a group of components, rather than any single component alone. In contrast, redundant interactions involve multiple components providing overlapping or duplicate information (such as strongly co-varying variables). Synergistic and redundant interactions between genes play crucial roles in biological systems, influencing phenotype and drug response. Studies have demonstrated synergistic interactions can help elucidate cooperative dynamics within gene networks and predict non-additive effects observed in drug combinations, while redundancy in gene copies can provide advantages under conditions promoting rapid growth and translation, highlighting the complex interplay between these types of interactions in genetic networks ^17,18^. One example of synergy is cooperativity, where system components (e.g., channel conductances, gene expression) influence excitability through interdependent, often multiplicative effects. While cooperativity can enhance sensitivity, it may also introduce fragility. In contrast, degeneracy describes the system’s capacity to achieve a similar pattern of activity through multiple distinct combinations of underlying components. This implies a high-dimensional solution manifold, where both synergistic and redundant interactions can coexist. Complementing this, degeneracy can also arise from partial functional redundancy among system components. Each of these modes of interaction (degeneracy, synergy, including cooperativity) supports robustness through diverse, distributed, and compensatory mechanisms.

The focus of our study is to investigate how high-dimensional interactions among biophysical model parameters and gene expression patterns relate to neuronal electrical phenotypes. We apply a unified information-theoretic framework to both conductance-based neuron models and single-cell transcriptomic data to quantify higher-order dependencies that shape excitability and firing behavior. Building on our previous work ^19,20^, we generate populations of excitatory and inhibitory cortical neuron models that replicate similar electrophysiological features despite varying in their underlying parameters. This enables us to assess how combinations of ionic conductances contribute to specific electrical properties. To quantify these dependencies, we apply Organisation information (O-info) ^21^ to assess global dependencies among parameters, and the Redundancy–Synergy Index (RSI) ^22^ to relate these interactions to specific electrophysiological features. Additionally, we analyze publicly available Patch-seq and single-cell mRNA sequencing data from cortical interneuron and pyramidal neuron populations ^12,23^ to investigate high-order structure in gene expression and compare it with the interaction patterns observed in model parameters. We hypothesize that the population of conductance-based models generated through stochastic sampling can serve as a “null model” for biological variability. In this framework, deviations from the “null model” may reflect biological constraints, such as transcriptional co-regulation of ion channels, or methodological factors, such as sampling strategy. This perspective underscores the complexity and variability of the relationships between gene expression and neuronal function, and highlights the importance of comparing observed structure against appropriately sampled generative baselines.

## Results

### Synergistic high-order interactions of ion currents in biophysically detailed models

To investigate the presence and impact of high-order interactions on the electrical phenotype (e-phenotype) of neurons, we used a detailed biophysical model of Layer 5 Pyramidal Cells (L5PC) of the somatosensory cortex (SSCx) (Fig. 1.A) ^19^. This model, incorporating region-specific distributions of ion channels and calcium dynamics across the soma, dendrites, and axon (present in the model as AIS followed by a myelinated compartment), underwent parameter exploration using a Markov Chain Monte Carlo (MCMC) approach ^20^. Through this process, we generated 110128 models capturing 60 electrical features of L5PC, presenting a wide range of ion channel conductances, calcium dynamics, and other biophysical properties. This process created a population of parameter sets representing a single e-phenotype of L5PC neuron. Importantly, this population of models display a variability in electrical features similar to the biological diversity observed in L5PC biological population ^20^.

**Figure 1.**
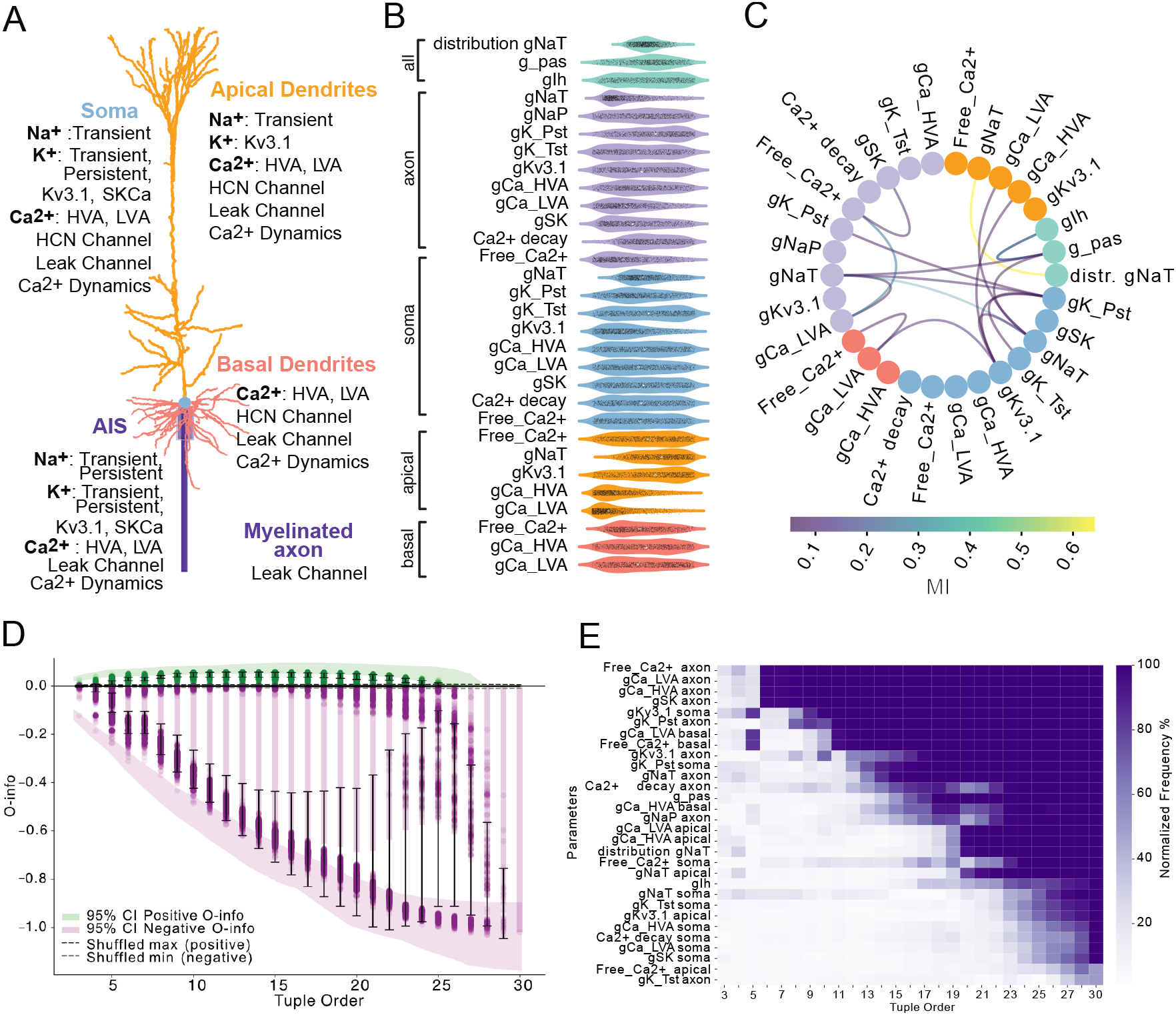
Analysis of a biophysically detailed L5PC neuron model and parameter interaction. **A.** Morphology of the L5PC neuron model, showing the distribution of ion channels across soma, dendrites, AIS, and axon. **B.** Parameter distributions from models generated using MCMC. **C.** MI map between parameter pairs. The colors of the nodes correspond to the color of the compartments of the neural morphology from A and B. The color of the edges corresponds to the MI value. **D.** O-info plot showing the relationship between tuple number (number of parameters) and O-info values. Each dot represents the O-info value for a specific tuple. The bars and whiskers illustrate the mean and standard deviation of O-info values at each tuple order. The color encodes positive (green) and negative (purple) O-info values. Shaded regions indicate the 95% confidence intervals computed via bootstrapping (50 repetitions, sample size = 1000). Dashed lines represent the maximum observed positive and negative O-info values obtained from 100 shuffled surrogates of the original data. **E.** Normalized frequency map of parameters frequently appearing in top 5% of high-synergy tuples.

The resulting parameter distributions revealed substantial variability in conductances such as gIh and gK Pst, as well as in the parameters reflecting calcium decay and free calcium levels across neuronal compartments (Fig. 1.B). This parameter variability underscores both variability in e-phenotype but also the degeneracy of the parameter space. Moreover, the parameter space was homogeneous and independent of the model fitness (see Fig. S1.A). Next, we examined the relationships between these parameters by calculating pairwise mutual information (MI), which revealed significant interdependencies (Fig. 1.C). Notable interactions included axonal sodium conductance (gNaT in axon) and its decay constant along the apical trunk, correlations between axonal and somatic sodium conductances, and strong interactions between calcium conductances (gCa HVA, gCa LVA) and intracellular calcium levels. Overall, 17 out of 30 parameters demonstrated statistically significant MI (see Methods section), indicating that the e-phenotype is supported by a degenerate solution space in which extensive parameter interdependencies emerge as signatures of the system’s flexibility.

To move beyond pairwise interactions, we applied high-order information measures to characterize the presence and nature of multi-parameter interactions ^21^. We first used O-information (O-info), a measure able to reveal redundant (O-info> 0) or synergistic (O-info< 0) interactions between variables. We computed O-info for combinations of 3 parameters, 4 parameters, and so forth up to 30 parameters, thereby spanning the full range of possible multi-parameter interactions (Fig.1D). The minimum of O-info values decreased consistently as the tuple size increased, reaching a minimum for 25 parameters. Beyond this point, adding more parameters did not further change the O-info measure. The negative value of O-info indicated a strong prevalence of synergistic interactions between model parameters. Synergistic interactions, in this context, are akin to collective computations within the model, where multiple parameters interact in a non-additive manner to create outcomes that would not be predictable by examining each parameter individually. These synergistic effects suggest that certain groups of ion channels and calcium dynamics act together to influence the models’ behavior in a way that goes beyond the sum of individual influences. Notably, the O-info associated with redundancy (O-info> 0) was minimal, approximately 10 times lower than that for synergy, highlighting the dominance of synergistic interactions (Fig.1.D). These patterns contrasted with those observed in the shuffled control ensembles, where column-wise randomization resulted in O-info values fluctuating around zero. Interestingly, when restricting the analysis to higher-cost L5PC models (optimization score > 7), we observed a marked increase in redundancy and a decrease in synergy values (Fig. S1.C), indicating that model quality influences the balance of high-order interactions.

To identify the specific parameters that contribute most significantly to the minimum O-info values, we analyzed the composition of parameter combinations associated with the highest levels of synergy. Specifically, we focused on the parameters that appeared most frequently in the top 5% of tuples with the lowest O-info values for each tuple order. For each tuple size (e.g., combinations of 3 parameters, 4 parameters, and so on), we calculated how often each parameter appeared in the most synergistic tuples, providing a frequency score that reflects each parameter’s contribution to high-synergy interactions across different tuple orders (Fig.1.E) (for the redundancy related map see Fig. S1.B). Key contributors to these synergistic interactions included parameters related to calcium dynamics, such as Free Ca2+ (in both soma and dendrites) and Ca2+ decay, which frequently appeared in the most synergistic tuples. Additionally, potassium channel conductances, specifically gK Pst and gKv3.1, were also frequently present in top synergistic tuples. This indicates that potassium conductance is another major factor shaping the collective dynamics, potentially influencing the model’s capacity to generate or shape its electrical features. Other channels, including gNaT, gCa LVA, and gCa HVA, showed consistent but moderate contributions to synergy. Interestingly, the parameter that appeared least frequently in the most synergistic tuples was axonal gK Tst. This aligns with our parameter sensitivity analysis ^19^, which showed that removing gK Tst from the model had minimal impact on the model’s output. This finding suggests that gK Tst plays a limited role in driving the simulated neuron model’s collective, synergy-based dynamics and may not be critical for the model’s overall behavior.

### Dependencies between electrical features and model parameters in biophysically detailed models

To map out the relationships between the electrical phenotype and the underlying ionic currents, we examined the relationship between the neuronal model outputs (electrical features, or “e-features”) and the model parameters. First, we extracted e-features from voltage traces produced by each generated model (Fig.2.A left). We then computed MI between each pair of e-feature and biophysical parameters to quantify the dependencies. All e-features showed statistically significant MI values, ranging from 0.01 to 1. For further analysis, we focused on e-feature/parameter pairs with MI greater than 0.1 (Fig. 2.A right).

**Figure 2.**
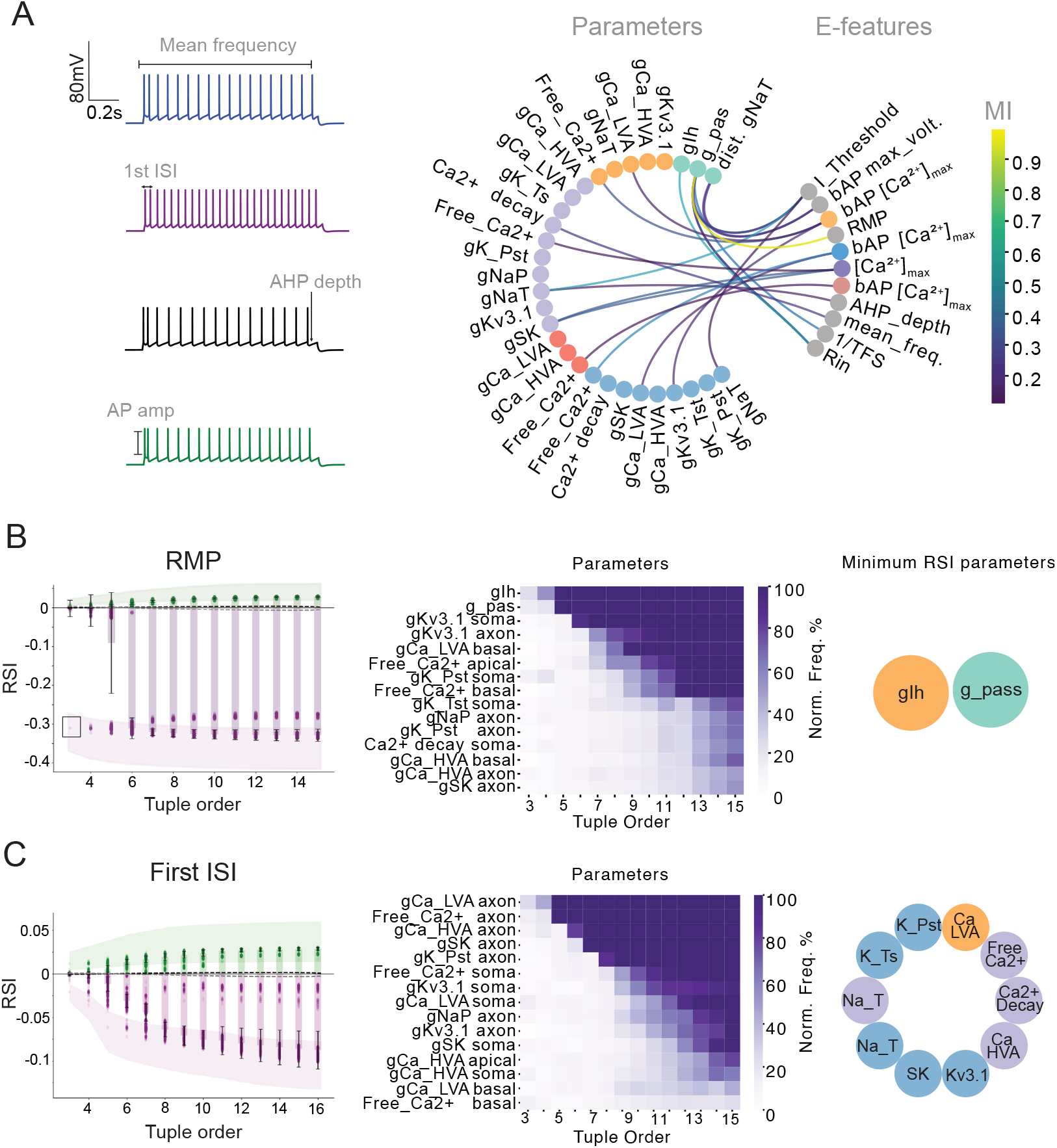
Pairwise and high-order interactions between electrical feature and model parameters in L5PC model. **A**. Left: Example voltage traces from generated L5PC models in response to a step current injection (200% of their rheobase), along with examples of the extracted electrical features. Right: MI map showing relationships between model parameters and e-features. Parameters are color-coded by compartment (Fig. 1.A); e-features are shown in gray, except for bAP features, which match their compartment color. **B.** Left: RSI values as a function of tuple order for RMP. Shaded regions show 95% confidence intervals from 50 bootstrap repetitions (sample size = 1000). Dashed lines mark the extrema from 100 shuffled surrogates. Middle: Parameters most frequently found in the top 5% of minimum-RSI tuples. Right: Schematic showing parameters with minimum RSI values, colored by anatomical location. C. Similar analysis as in B, but for the First ISI (Interspike Interval) feature.

High MI values revealed distinct relationships between e-features and specific parameters. Calcium concentrations in the axon and back-propagating action potential (bAP) features in the basal apical dendrites displayed strong associations with calcium-related parameters in the respective compartments. Somatic and axonal sodium conductances each exhibited dependencies with rheobase (I Threshold), while action potential (AP) amplitude and afterhyperpolarization (AHP) depth were influenced by axonal sodium conductances. Mean spiking frequency showed a strong relationship with axonal calcium decay, whereas passive conductances, including g pas, shared high MI with the time to first spike (the feature used is the inverse of time to first spike 1/TFS). Input resistance (Rin) and resting membrane potential (RMP) both displayed high MI with g pas (Fig. 2.A right).

In order to determine whether specific sets of parameters collectively determine specific e-features, we moved beyond pairwise interactions and explored higher-order interactions between parameters and specific e-features. We focused on two e-features: resting membrane potential (RMP) and the first interspike interval (ISI). Intuitively, these e-features are expected to involve distinct parameter sets due to their different nature: RMP should mainly rely on passive conductances while ISI is influenced by active conductances. Additionally, while RMP shared high pairwise MI with two distinct parameters, the first ISI did not have any statistically significant MI with individual parameters. To quantify higher-order information between parameter sets in relation to each e-feature, we applied the Redundancy-Synergy Index (RSI) ^22^, which allowed us to compute conditional mutual information and therefore understand which parameter interactions were responsible for a given e-feature. As for O-info, negative RSI values indicate synergistic interactions, while positive values reflect redundant interactions. For RMP, the RSI reached its lowest value (−0.35) with just two parameters. With additional parameters, the RSI did not increase, remaining at −0.35, suggesting that any additional parameters do not contribute significantly to RMP’s RSI (Fig. 2.B left). When examining the identity of these parameters, we found that they are gIh and g pas (Fig. 2.B middle, right). This aligns with findings in pyramidal cells that demonstrated the implication of Ih and passive current in shaping somatic RMP ^24^. These parameters formed a synergistic module specifically for RMP, indicating that their joint variations, rather than individual effects, were critical for regulating this feature. While not implying a transcriptionally co-regulated module in the genomic sense, this synergy reflects a functional coordination at the level of conductance tuning—potentially emerging through homeostatic plasticity mechanisms that respond to phenotype-level error signals. In contrast, when examining higher-order information for the first ISI, we observed that RSI values continued to decrease as the tuple size increased, stabilizing around 10 parameters (Fig. 2.C left). The parameters constituting the maximum synergy for ISI included axonal calcium and sodium conductances, somatic potassium and sodium conductances, and apical calcium conductances (Fig. 2.C middle, right). This result suggests that ISI is influenced by a more complex module involving multiple conductances across various compartments.

Overall, these results demonstrate that the behavior of the L5PC neuron model is governed by complex, synergistic interactions among parameters, particularly those related to calcium handling and potassium conductance across different compartments. These synergistic modules are specific to certain e-features, and their complexity can be revealed by examining higher-order interactions. These findings underscore the importance of multi-parameter synergy in shaping distinct aspects of neuronal function, with RMP and first ISI being controlled by distinct, high-order parameter modules.

### High-order interactions are specific to the electrical phenotype

So far, we have examined the high-order interaction modules for a single electrical phenotype, the L5PC neuron. However, an open question remains: would these high-order interaction modules differ in models producing different electrical phenotypes? To investigate this, we examined two distinct electrical models of cortical interneurons from the somatosensory cortex (SSC): the continuous accommodating cells (cAC) and the bursting non-accommodating cells (bNAC) ^25^. Both models were optimized to match the respective electrophysiological recordings obtained from these neuronal types ^19^ and share identical parameter sets and morphology, allowing for a meaningful comparison of optimized parameter values (Fig. 3.A).

**Figure 3.**
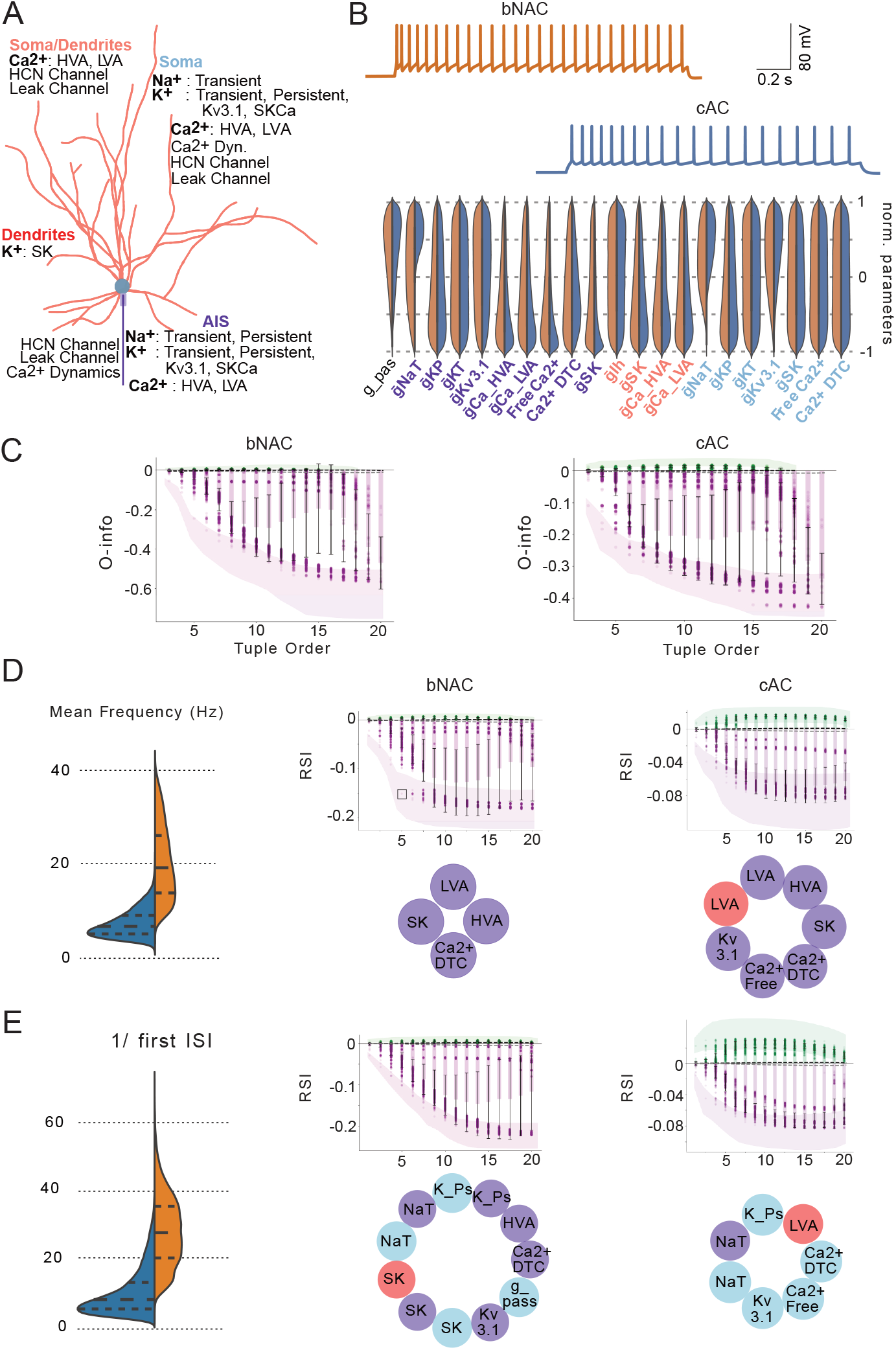
High-order interaction in the models of bNAC and cAC interneurons. **A.** Morphology of the interneuron model used for optimization, illustrating the distribution of ion channels across the soma, dendrites, AIS, and axon used in the model. **B.** Top: Example voltage traces from the continuous accommodating (cAC) and bursting non-accommodating (bNAC) models (in response to IDrest protocol, 150%/140% from the rheobase). Bottom: Distribution of optimized parameters, with values for cAC shown in blue and bNAC in orange. **C.** O-info values as a function of tuple order, showing the progression of high-order interactions for bNAC (right) and cAC (left) models. Shaded regions indicate the 95% confidence intervals computed via bootstrapping. Dashed lines represent extrema values of O-info for shuffled data. **D.** Left: Mean firing rate distribution for bNAC (orange) and cAC (blue). Middle: RSI plot for bNAC, showing the minimum parameter set driving high synergy (with colors corresponding to their anatomical location in the neuron model). Right: RSI plot and minimum parameter set for cAC, similar to the middle panel. **E.** Similar analysis as in D for the inverse First ISI feature.

When we compared the distributions of values of optimized parameters between the two models, we found that 19 out of 21 parameters exhibited similar distributions. The only significant differences were observed in the axonal and somatic sodium conductances (gNaT), which varied between the two models (Fig. 3.B). This indicates a high degree of similarity in the underlying parameter distributions between the cAC and bNAC models, with minimal differences confined to sodium conductances.

Furthermore, both models displayed a predominance of synergistic interactions across parameter tuples, as evidenced by the consistently negative O-info values. For both models, the O-info values stabilized at a tuple order of 14, indicating that beyond this point, additional parameters do not contribute significantly to the synergy or redundancy in the model (Fig. 3.C). This stabilization suggests that a similar number of parameters is required to capture the full scope of synergistic interactions in both electrical phenotypes (see Fig. S2). We also observed that the range of O-info values differs between bNAC and cAC model populations, with bNAC models generally exhibiting stronger synergy. This difference likely reflects distinct high-order parameter interactions or dynamical constraints specific to each cell type, suggesting that the two model classes may rely on qualitatively different computational strategies.

Given the similarity in the initial parameter space, we explored whether the same modules of biophysical parameters could drive electrical features in each model. Specifically, we focused on the features reflecting the firing pattern of the cells: mean firing frequency and inverse first ISI (1/ISI) features in response to a current injection of 200% of the rheobase. Notably, the mean firing frequency for the bNAC model was 20 Hz (± 7.6), higher than the mean frequency of 7.4 Hz (± 2.9) for the cAC model (Fig. 3.D). The RSI values for both models indicated strong synergy. For the bNAC model, the minimum tuple order that achieved the most negative RSI (indicating maximum synergy) was 5 (4 parameters), whereas for the cAC model, it was 8 (7 parameters). In the bNAC model, axonal LVA and HVA calcium channels, SK channels, and calcium dynamics formed the minimum parameter set that produced maximum synergy. For the cAC model, the minimum synergistic module included the same parameters as for bNAC, along with somatodendritic LVA calcium channels, axonal Kv3.1 channels, and free calcium. Interestingly, at a tuple order of 6, both models shared the same core parameters, indicating close similarities between their underlying interaction structures.

To examine whether this similarity extends to other features, we tested whether the co-variation modules for bNAC would be a subset of those for cAC for the inverse first interspike interval (ISI). Consistent with their electrical phenotypes, the inverse first ISI was larger for bNAC (29.9 ± 10.3) than for cAC (9.6 ± 6.7) (Fig. 3.E, right). For the bNAC model, we observed a steep and prolonged decay in RSI values, requiring 12 parameters to reach the minimum RSI, indicating significant synergy (Fig. 3.E, middle). In contrast, only 7 parameters were needed to achieve minimal RSI values in the cAC model (Fig. 3.E, left). Notably, only a quarter of the parameter sets overlapped between bNAC and cAC for this feature, indicating that the biophysical modules are highly specific to the electrical phenotype.

In summary, cAC and bNAC models, built from the same parameter space and morphology, exhibit nearly identical parameter distributions. Both display a predominance of synergy, highlighting the importance of cooperative parameter interactions in shaping distinct firing phenotypes. While cAC and bNAC models share similarities in their synergistic parameter modules for certain e-features, these modules are distinct and phenotype-specific for other e-features. The specificity of biophysical modules defining specific e-features highlights that the complexity of electrical phenotypes emerges from unique sets of synergistic interactions, tailored to the firing behavior characteristic of each neuronal type.

### Ion channel mRNAs as key elements identifying neuronal types

To investigate high-order interactions beyond in silico model populations, we leveraged patch-seq data ^12^ to explore the relationship between mRNA expression levels and electrical features in biological cortical GABAergic interneurons. Since our in silico results focused on interactions between ion currents, our first step was to determine the role of ion channel expression levels in defining principal interneuron subtypes.

We began by isolating ion channel genes (a set of 150 among 45 768 mRNA) from the gene expression data ^12^ and performed clustering, aiming to capture the main interneuron types. When we compared our clustering results with the ground truth classifications provided by the dataset (Fig. 4.A), we found that most primary clusters were well-replicated using only these 150 ion channel genes. Notably, we observed distinct clusters for PV, SST, and LAMP5, indicating that ion channel gene expression alone effectively identifies these major interneuron subtypes. However, the clustering of ion channel genes merged SNCG and VIP into a single cluster, and an additional two clusters emerged—one corresponding to a mixed population of the primary cell types and the other to a distinct SST subtype.

**Figure 4.**
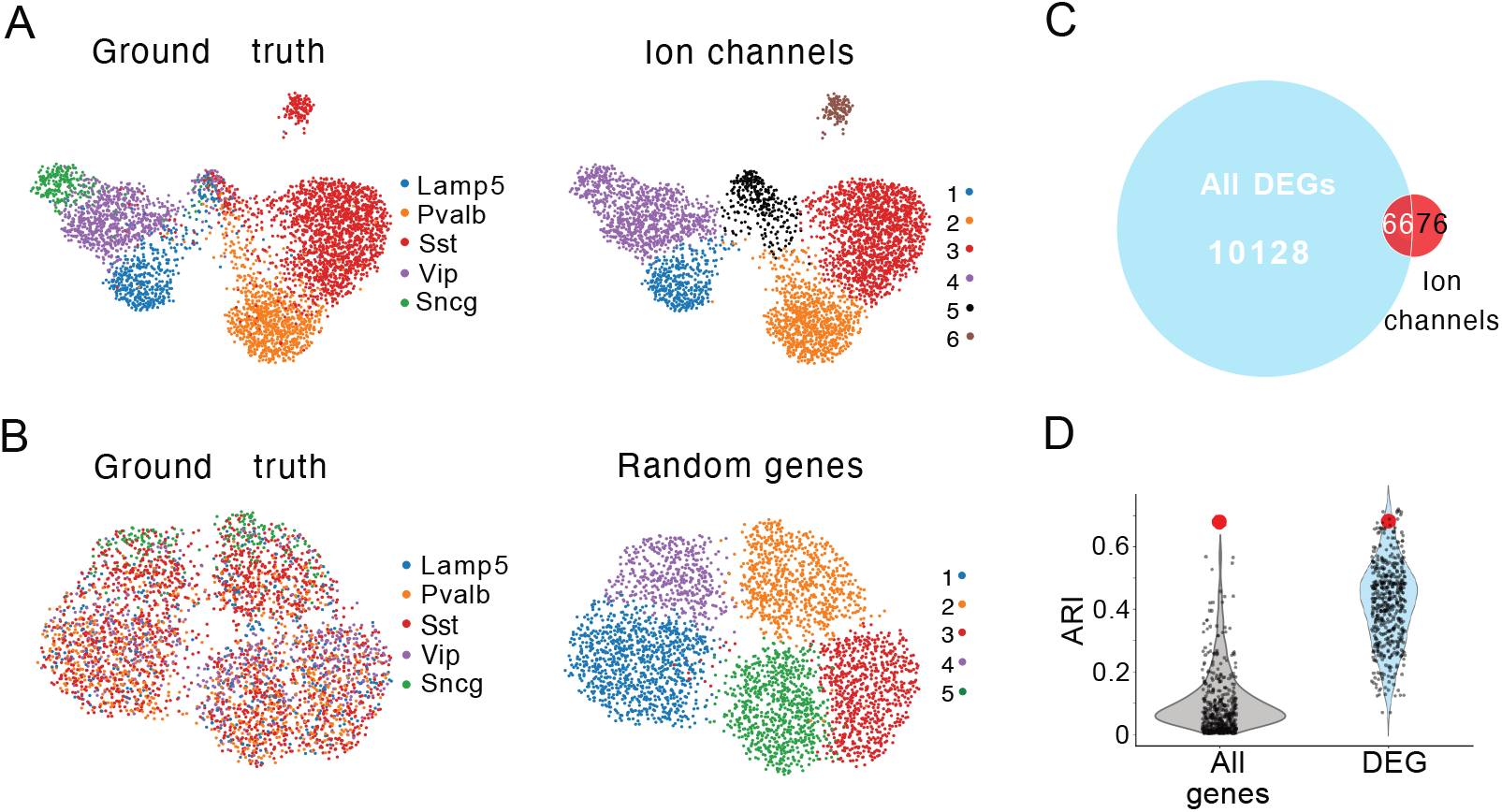
Transcriptomic clustering of interneurons using Ion channel genes vs. random gene sets and differential expression analysis. **A.** UMAP representation of IN transcriptomic clustering based on ion channel-encoding genes. Each point represents single cell. Left: clusters labeled according to ground truth cell types for comparison. Right: same UMAP, with clusters identified by ion channel gene expression, each cell is labeled with a unique identifier. **B.** UMAP representation of interneuron transcriptomic clustering based on a randomly selected gene subset. Left: cells labeled according to ground truth cell types. Right: cells labeled according to clustering results.. **C.** Venn diagram showing the overlap of DEGs identified through ground truth clustering and ion channel encoding genes. **D.** Distribution of the ARI values for clustering based on randomly selected genes. Grey distribution represents ARI values for clustering with random genes sampled from the entire gene pool, while blue distribution represents ARI values for clustering with random genes from the DEGs. Each dot corresponds to a single random sampling of the corresponding condition. The ARI value for clustering based on ion channel-encoding genes is marked by a red dot.

To test whether this clustering was specific to ion channel genes, we repeated the clustering process with randomly selected mRNA subsets. We randomly sampled 150 genes 500 times and used these samples for clustering (Fig. 4.B), comparing the Adjusted Rand Index (ARI) of the resulting clusters with the ground truth (Fig. 4.C). The ARI of the ion channel gene clustering was 0.69, at least seven times higher than the average ARI obtained from random gene sets (mean ARI of 0.05; grey violin plot) (Fig. 4.D). This difference suggests that ion channel genes have substantial predictive power for interneuron subtypes compared to random mRNA subsets.

An additional validation was performed by examining differentially expressed genes (DEGs) (see Methods) to control if a more specific subset of mRNAs could match the clustering power of ion channel genes. Using the ground truth, we identified DEGs across cell types and noticed that only a subset of ion channel mRNAs was comprised in these DEGs (Fig. 4.C). We then sampled 150 DEGs 500 times and performed clustering. The average ARI for DEGs (mean ARI of 0.4; blue violin plot) was higher than the ARI for randomly sampled genes from the full gene pool (mean ARI of 0.05), showing that DEGs generally have better clustering performance. Importantly, the ARI for ion channel genes was within the top 1% of the ARI distribution generated by DEG clusters, confirming that ion channel genes have a predictive power greater than 99% of DEG subsets (Fig. 4.D). In summary, these findings demonstrate that ion channel gene expression alone provides a robust basis for distinguishing interneuron subtypes, outperforming both random mRNA samples and most DEG subsets. This result emphasizes the unique and predictive role of ion channels in defining neuronal identity and capturing the main structural and functional variations among interneurons.

### High-order interactions between ion channel mRNAs specific to the electrical phenotype

To investigate whether ion channel genes share statistical interactions specific to neuronal subtypes, we focused on two electrically different interneuron types: Lamp5 and Pvalb. Pvalb interneurons are characterized by a fast-spiking pattern of activity, while Lamp5 neurons exhibit slower, delayed spiking ^12^. We selected 50 ion channel genes with non-zero median expression in both subtypes, including 2 HCN channels, 6 calcium channels and subunits, 6 chloride channels and subunits, 4 sodium channels, and 32 potassium channels.

First, we analyzed pairwise interactions between these ion channel genes using mutual information (Fig. 5.A). The Lamp5 neurons exhibited higher MI values, reaching up to 0.26, compared to Pvalb neurons, which peaked at 0.19. In total, 58% of the ion channels in Lamp5 displayed statistically significant pairwise interactions, whereas only 34% of ion channels in Pvalb did so. In Lamp5 neurons, these interactions involved a broad range of potassium channels (Kcnq, Kcnk, Kcnj, Kcnt, Kcnc, Kcnn, and Kncd families), as well as calcium channels (Cacna1 family), chloride channels (Clcn family), and sodium channels (Scn family). In contrast, the Pvalb neurons exhibited a more focused pattern, with strong correlations primarily among potassium channels from the Kcn family, along with select sodium channels from the Scn family.

**Figure 5.**
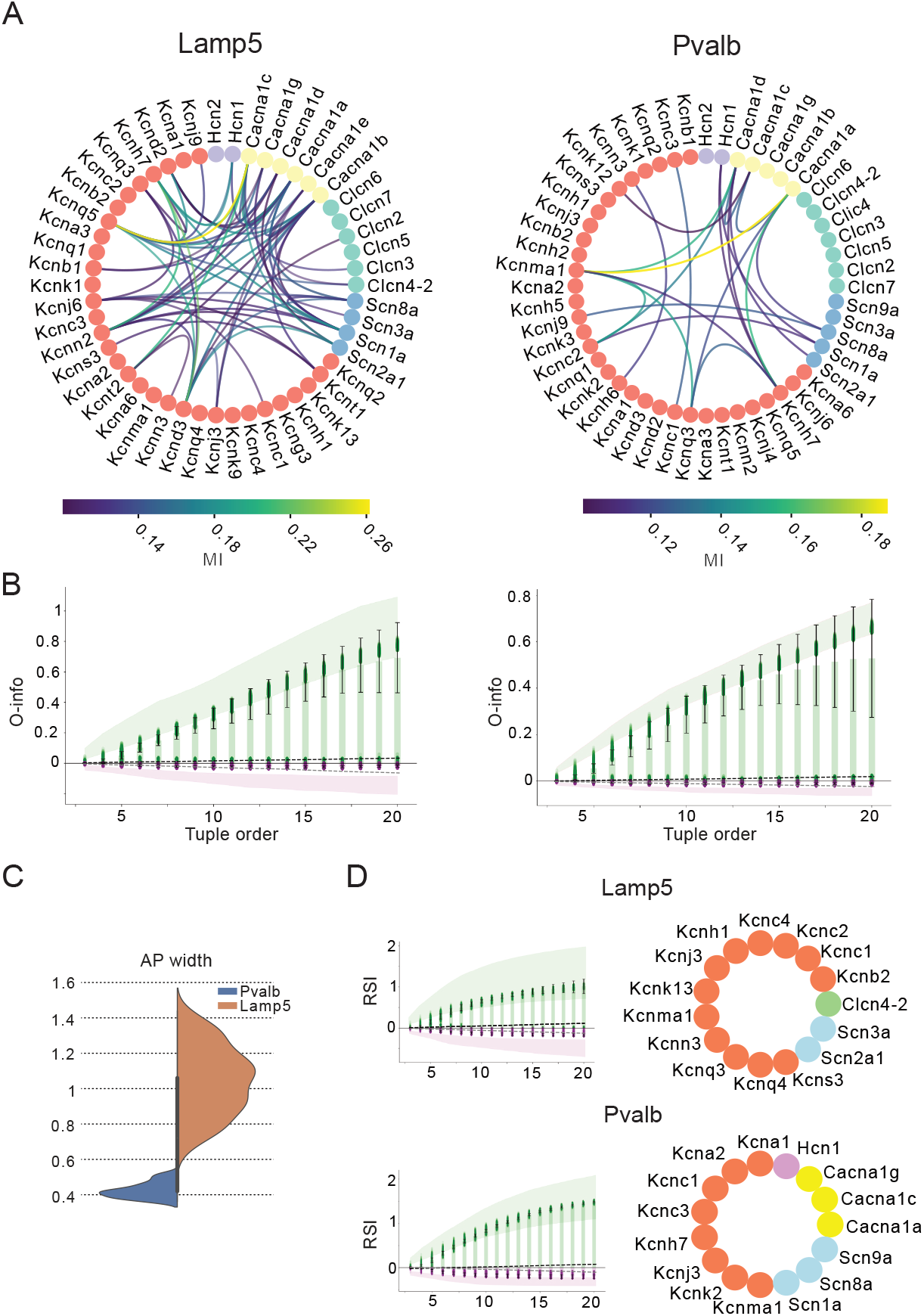
High order interactions in the Ion Gene expression data. **A.** A map of 50 ion channel-encoding genes. Node color represents the general current type: potassium (coral), sodium (blue), chloride (green), calcium (yellow), and HCN (purple) families. The color bar indicates the MI values. Left: MI map for Lamp5 ion channel gene expression. Right: same for Pvalb. **B**. O-info values plotted as a function of tuple order for Lamp5 (left) and Pvalb (right) neurons. Shaded regions indicate the 95% confidence intervals computed via bootstrapping. Dashed lines represent extrema values of O-info for shuffled data. **C**. Distribution of action potential (AP) widths (in milliseconds) for cells from Lamp5 and Pvalb neuron types. **D**. RSI values as a function of tuple number (right) and corresponding set of the genes that result in highest value of RSI (left) for Lamp5 (top) and Pvalb (bottom) neurons.

We then examined high-order interactions among ion channel genes for both cell types. Both cell types showed a strong prevalence of redundant interactions (positive O-info), as indicated by negative O-information values fluctuating around zero. The O-info increased monotonically with tuple size, with potassium-encoding genes dominating the tuples that exhibited the highest redundancy (see Fig. S3).

To explore the link between electrical features and ion channel gene expression, we analyzed electrical features from patch-seq data alongside gene expression profiles from the same individual cells for both Lamp5 and Pvalb neurons. We focused on features that distinguish these cell types, particularly action potential half-width (AP half-width), known to be shorter in Pvalb neurons (thus allowing fast-spiking). The AP width in Pvalb was threefold shorter (0.4± 0.1 ms) than in Lamp5 (1.1 ± 0.5 ms). Redundancy-Synergy Index (RSI) analysis again revealed predominantly redundant interactions among ion channel genes in both types. The tuple order yielding the highest RSI was similar between cell types (15 for Lamp5 and 14 for Pvalb). However, the genes driving maximal RSI for AP width differed: in Lamp5, potassium channel genes predominated, along with sodium channels and one chloride channel, while in Pvalb, we observed contributions from Hcn1, multiple calcium channels, and sodium channels. We further extended our analysis on the rheobase feature and similarly found implication of distinct co-expression modules (see Fig. S4 and S5).

These findings suggest that, although both cell types exhibit redundant high-order gene interactions, the specific ion channel genes contributing to these interactions differ between Lamp5 and Pvalb, reflecting their distinct electrophysiological properties. Moreover, while in silico populations displayed mostly synergistic interactions between ion conductances, biological populations displayed strong redundant interactions between ion channel genes.

### Bridging the gap between the Models and Transcriptomics results

While our initial analyses revealed a dominance of synergistic interactions in uniformly sampled conductance-based models, and a contrasting dominance of redundancy in gene expression data, the origin of this discrepancy remained unclear. Thus we investigated whether this observed discrepancy can be explained by the covariance structure or by the sampling strategy of the parameter space. To address this, we employed two complementary strategies: (1) reanalyzing a dataset of biophysical models fitted to pyramidal neurons using a different sampling strategy ^26^, and (2) subsampling high-covariance models from a uniformly sampled interneuron population.

In the first approach, we asked whether a shift toward redundancy could be observed in models generated through an entirely different sampling strategy, specifically, one that reflects targeted optimization around biological data rather than stochastic exploration. To this end, we analyzed a previously published dataset of conductance-based models fitted to L4/L5 pyramidal neurons from the mouse visual cortex using a multiobjective evolutionary algorithm (IBEA) ^26,27^. Although these models were not generated in the present study, they offer a valuable point of comparison, as they exemplify a data-constrained, non-random modeling approach in which multiple solutions are clustered around fixed electrophysiological targets.

To ensure comparability, we selected only those neurons whose electrophysiological properties were tightly clustered into a single electrical type (as classified in ^28^), yielding a subset of 72 neurons. For each of these neurons, a population of 40 models had been generated, each representing a distinct solution from the final generation of the evolutionary optimization, known as the Pareto-optimal “Hall of Fame” (HOF) set. These 40 solutions per neuron are all consistent with the target electrophysiological features, but differ in their underlying ion channel parameters.

We first analyzed the full set of 2,880 HOF models (72 neurons × 40 models) and computed O-information across increasing tuple orders. Despite all models matching their electrophysiological targets, the interaction profiles exhibited a clear emergence of both redundancy and synergy, with redundancy dominating at higher tuple orders (Fig. 6.A).

**Figure 6.**
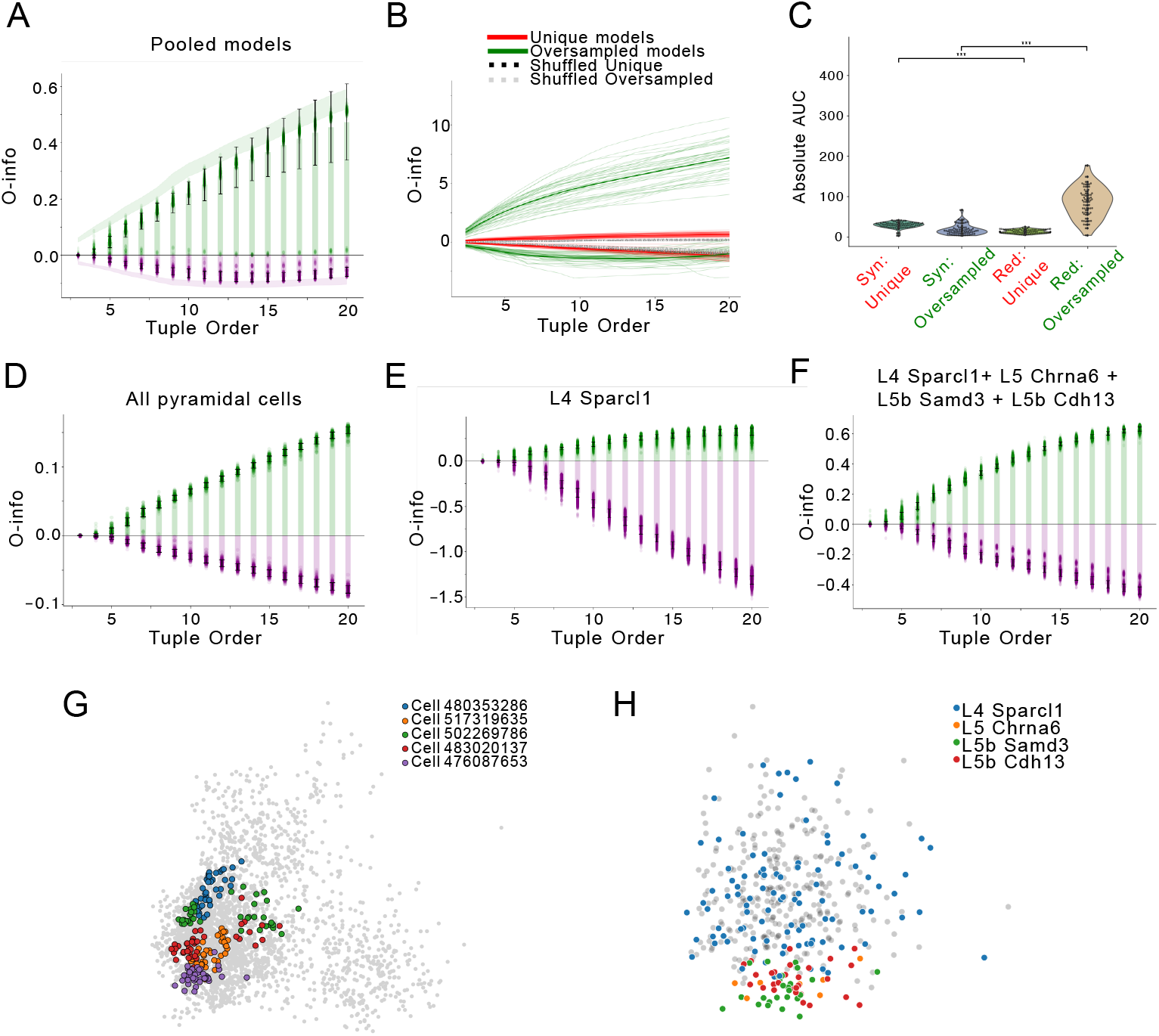
Model parameter and gene expression space shapes high-order interactions. **A.** O-info across tuple orders for all 2,880 models. Green (positive) and purple (negative) values indicate redundancy- or synergy-dominated interactions, respectively; lines show mean ± SD. Shaded areas denote 95% confidence intervals from bootstrapping. **B.** O-info for two model groups: “Unique” (72 models from distinct HOF sets) and “Oversampled” (72 models from two cells, 36 each). Gray and blue dashed lines represent control groups with parameter-wise shuffled models. **C.** Distribution of absolute O-info AUC values (minima for synergy, maxima for redundancy) in Unique vs Oversampled groups. Violin plots show variability; Mann–Whitney U test, p ¡ 0.001. **D.** O-info across tuple orders for all L4/5 pyramidal neuron models from VISp. **E.** O-info across tuple orders restricted to models from the L4 Sparcl1+; curves as in (D). **F.**O-info across tuple order for a combined ensemble of four excitatory subtypes (L4 Sparcl1+, L5 Chrna6+, L5 Samd3+, and L5 Cdh13+). **G.** PCA projection of the 2,880 models’ parameter space (from panel A), with five randomly selected cells highlighted in color (one dot = one HOF model). The gray background shows the distribution of models from other models. **H.** PCA projection of the ion gene expression of the cells shown in panel D. Each dot represents one cell, with color indicating its excitatory subtype cluster.

We next examined how sampling strategy affects the prevalence of synergy and redundancy. Specifically, we compared two equally sized subsets of 72 models: (1) a “Unique group” comprising one randomly selected HOF model from each of the 72 neurons, and (2) an “Oversampled group” comprising 36 models from each of two neurons. The Unique group exhibited synergy-dominated interaction profiles, whereas the Oversampled group showed pronounced redundancy (Fig.6.B). This effect was not due to sample size alone: both groups contained the same number of models. Rather, it reflected the structure of sampling: the Unique group combined models from diverse clusters, forming a broad distribution, whereas the Oversampled group isolated two tight clusters, where constrained parameter relationships drove redundancy. Dimensionality reduction of the full model ensemble further illustrates this point: PCA reveals that models from the same optimization run (i.e., 40 HOF models per cell) form tight, cell-specific clusters, while models from different cells are more broadly dispersed (Fig. 6.G).

To quantify these differences, we computed the absolute area under the curve (AUC) for O-info trajectories across tuple orders. The Unique group showed significantly higher synergy AUC than redundancy (p < 0.001), while the Oversampled group showed the opposite pattern (p < 0.001) (Fig. 6.C). Both groups significantly differed from their respective shuffled controls (Fig.S6.C), confirming that the observed effects were not due to sampling noise. These findings suggest that oversampling around a fixed biological target induces a redundancy-dominated structure, while sampling across diverse neurons allows synergy to emerge.

We then asked whether similar principles apply to biological gene expression. Using single-cell transcriptomic data from the mouse visual cortex ^23^, we analyzed 466 excitatory neurons from layers 4 and 5 and focused on ion channel–encoding genes. O-information analysis of expression profiles revealed a clear dominance of redundancy across all interaction orders (Fig. 6.D), closely paralleling the redundancy observed in oversampled biophysical model populations. Although these transcriptomic data originate from a separate study and are not directly linked to the modeled neurons, they sample from the same cortical layers (L4/L5 VISp) and offer a meaningful point of comparison.

To probe subtype-specific differences, we examined four well-defined excitatory neuron subtypes: L4 Sparcl1+ (n=75), L5 Chrna6+ (n=7), L5 Samd3+ (n=25), and L5 Cdh13+ (n=28). When analyzing the L4 Sparcl1+ subtype alone, we found a surprisingly strong synergy signature (Fig. 6.E), suggesting that within-subtype variability may encode complementary, high-order information. However, when we combined the L4 and L5 subtypes into a pooled ensemble, redundancy became dominant (Fig. 6.F). This shift echoes the pattern observed in oversampled model sets and suggests that mixing neuronal subtypes can dilute synergistic interactions and promote redundancy. It is further supported by the low dimensional projections of the combined transcriptomic dataset showing partial clustering by subtype (Fig. 6.H), confirming that the underlying gene expression distributions retain subtype-specific structure even after pooling. Notably, this effect was not observed when we combined two closely related L4 subtypes (Sparcl1+ and Scnn1a+ (n=95)), which appear intermixed in low-dimensional space; in this case, synergy remained slightly dominant rather than giving way to redundancy (Fig. S6.A,B).

Together, these results demonstrate that both synergy and redundancy can emerge in biophysical and biological systems, depending on how sampling captures the structure of underlying variability. Redundancy prevails when sampling isolates tightly clustered solutions—whether through repeated optimization around a fixed target or by focusing on biological subtypes. In contrast, synergy emerges when sampling spans across distinct clusters, revealing broader, less constrained parameter relationships. To further disentangle the origins of these high-order interactions, we next returned to our uniformly sampled cAC interneuron models, which previously exhibited strong synergy. Here, we asked whether it would be possible to reveal redundancy within this synergy-rich population purely by selecting models with high inter-parameter covariance. From the cAC models population, we selected a subset of models characterized by high inter-parameter covariance, using PCA and a covariance-based scoring metric (see Methods). Importantly, this selection preserved both the electrophysiological targets and the model structure—only the statistical dependencies among parameters were altered.

Strikingly, despite being drawn from a synergy-rich ensemble, the high-covariance subset showed a clear shift toward redundancy (Fig. 7.A, middle row), in stark contrast to equivalently sized random subsets, which retained synergistic interactions (Fig. 7.A, top row). This divergence was evident both in the O-info trajectories and in the MI matrices (Fig. 7.B): while the random subset showed minimal pairwise dependencies (Fig. 7.B, top row), the high-covariance subset revealed pronounced off-diagonal structure (Fig. 7.B, middle row), indicative of widespread parameter co-variations. A similar pattern of prevalence of the off-diagonal structure can be observed in the MI matrix of the ionic gene expression of Lamp5 (Fig. 7. B, bottom row).

**Figure 7.**
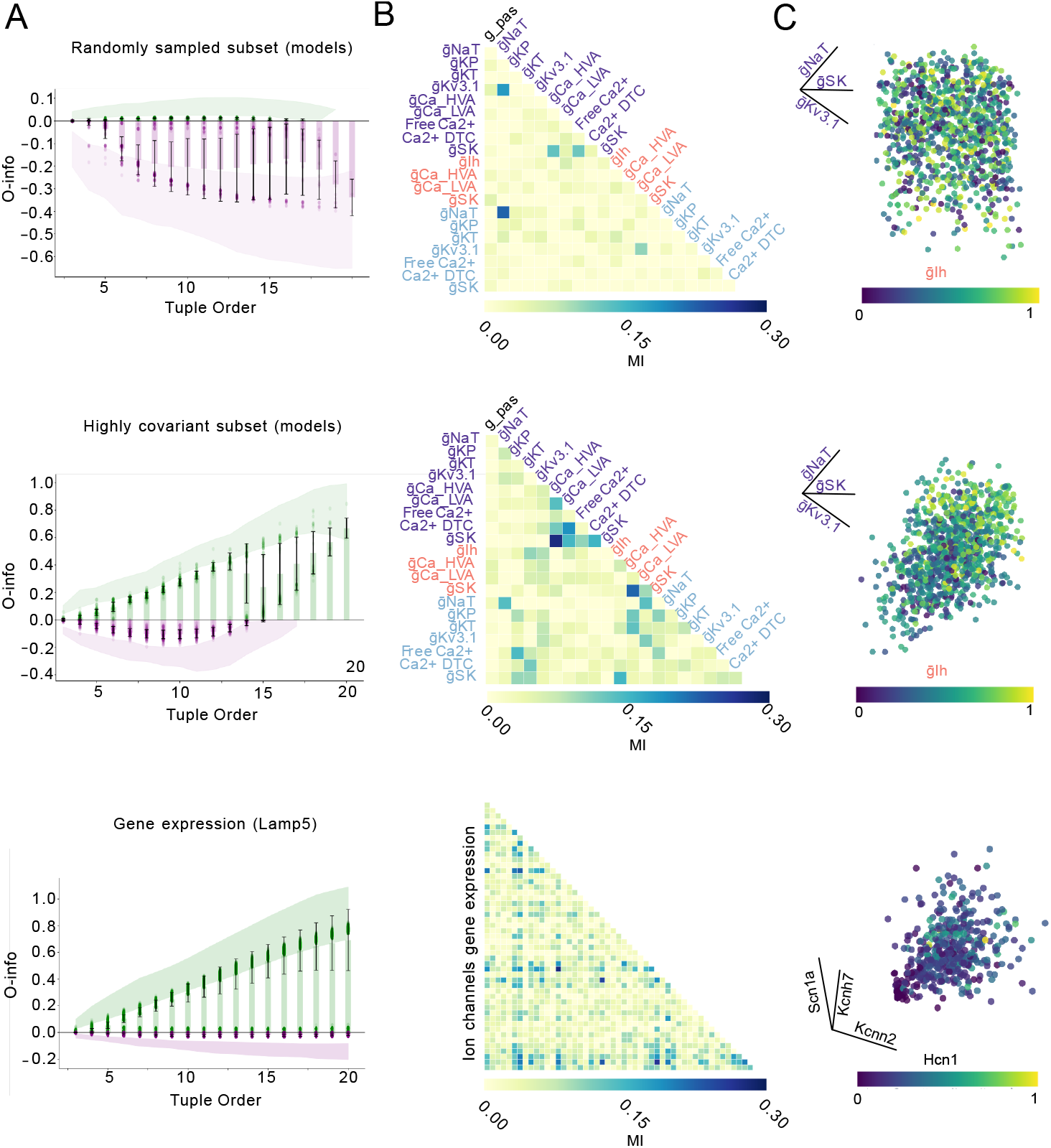
Covariance structure and the balance between synergy and redundancy in biophysical models and gene expression. **A.** O-info trajectories across tuple orders for three distinct datasets: a randomly sampled (n=1000) cAC model subset (top), a highly covariant model cAC subset (middle, n=1000), and Lamp5 gene expression data (bottom). Positive values (green) indicate redundancy-dominated interactions; negative values (purple) indicate synergy. Shaded regions show the full bootstrap range (n=50 repetitions of 100 samples each for cAC models, repetitions of 240 for Lamp5), and lines with error bars denote the mean ± s.e.m. **B.** Pairwise mutual information (MI) matrices for the same parameter or gene sets shown in (A). Only the lower triangle is shown, with parameter names rotated and placed along the diagonal. Color of the parameter names correspond to their anatomical location in the neuron model as in (Fig.3.A). **C.** Example 4D scatterplots highlighting relationships among four parameters or genes from each subset. In the top two panels (model subsets), the same set of four parameters is shown (e.g., gSK, gNaT, gKv3.1, glh), with color representing one dimension. In the bottom panel (gene expression), four ion channel genes (e.g., Hcn1, Kcnn2, Scn1a, Kcnh7) are shown for a Lamp5 interneuron population. All values were normalized and vary from 0 to 1.

To visualize these dependencies, we represented the 4D relationship among four example parameters (gNaT, gSk, gKv3.1,gIh) within each group (Fig. 7.C). In the randomly sampled subset, the joint distribution appeared diffuse and unstructured, consistent with high-dimensional independence. In contrast, the high-covariance subset revealed a clear geometric organization, suggestive of tightly coupled parameter relationships. Interestingly, this structured pattern was very similar to the one observed in gene expression data from Lamp5 interneurons (Fig. 7.C, bottom), where a selected set of ion channel genes (Scn1a, Kcnh7, Kcnn2,Hcn1) showed clear interdependence in both the MI matrix and the 4D projection. Together, these results reinforce a key insight: that redundancy emerges when parameters—or genes—become statistically entangled, whether through constrained biological regulation or methodological oversampling. Conversely, synergy is most prominent in diverse or decorrelated ensembles. This parallel between the biophysical and transcriptomic domains suggests that the observed interaction structure reflects population-level statistical organization, rather than intrinsic properties of the individual components.

## Discussion

This study explores the connection between biophysical parameters, gene expression, and neuronal electrophysiological phenotypes. Using biophysical models, statistical tools, and information theory, we examined how ionic currents shape neuronal behavior and linked these insights to transcriptomic profiles. We generated diverse electrical models via MCMC sampling, enabling systematic analysis of how combinations of ion channel parameters influence neuronal features. Our findings show that these features rarely depend on a single current, but instead emerge from complex interactions among multiple channels. This highlights the need for multivariate tools, such as O-info and RSI, to reveal the integrated effects of conductance interactions. Furthermore, our analysis of transcriptomic data identified gene expression modules associated with specific interneuron types, reinforcing the link between molecular and electrical identity. The prevalence of high-order redundancy among ion channel genes suggests a mechanism for robustness, degeneracy, and functional compensation. Potassium channel genes were especially dominant in redundant tuples, highlighting their role in stabilizing neuronal excitability. Notably, the specific genes driving maximal redundancy differed between Lamp5 and Pvalb interneurons, reflecting their distinct electrophysiological properties. This difference was particularly evident in the action potential width, a key distinguishing feature between these two interneuron types. Pvalb neurons are characterized by significantly shorter AP widths and lower rheobase values compared to Lamp5 neurons, aligning with their fast-spiking capabilities and higher excitability ^29,30^. The AP width influences the timing precision and frequency of neuronal firing, affecting synaptic transmission and the synchronization of neural networks ^31^. A narrower AP width in Pvalb neurons allows for rapid and high-frequency firing, which is essential for their role in controlling the timing of cortical circuitry ^29^. The shorter action potential width in Pvalb neurons, indicative of their fast-spiking nature, can be attributed to the specific combination of ion channels contributing to the maximal redundancy in this cell type. It is crucial to note that mRNA expression does not directly translate into protein expression ^32^. Future studies incorporating proteomic data would provide a more direct link between gene expression and functional ion channels.

The synergistic nature of the models and the redundancy observed in gene expression patterns raise interesting questions about the evolutionary advantages of these arrangements. Redundant elements in gene expression may provide robustness to the system, allowing for functional compensation if one component fails. In E.coli it was shown that the presence of multiple rRNA/tRNA gene copies can be advantageous under conditions that promote rapid growth and translation ^33^. This redundancy allows organisms to adapt more quickly to favorable environmental conditions ^33^. At the same time, redundancy offers significant advantages for both resilience and evolution. It has been shown that biological redundancy in protein-protein interaction networks enables robustness and increased tolerance for perturbations, enhancing their ability to maintain functionality despite genetic mutations or environmental stresses ^34,35^. Conversely, synergetic interactions in the model might offer more efficient information processing but at the cost of increased vulnerability if a critical component is disrupted.

In this study we directly addressed the possible causes of redundancy and synergy in both models and biological neurons, and provide answers about the discrepancies observed in these two datasets: while synergy between biophysical parameters dominated in most models, redundancy in ion channel expression dominated in most biological neuronal types. We show that this balance is shaped by inter-parameter covariance: even in broadly sampled MCMC models, selecting high-covariance subsets was sufficient to shift interactions toward redundancy. This result demonstrates that parameter co-variation alone, independent of biological constraints or fitting procedures, can suppress synergy by effectively reducing the dimensionality of the solution space. One possible interpretation is that biological solutions occupy a covariation-constrained subspace of a larger computationally adequate parameter space. The existence of redundancy-dominated subsets of models (see Fig. 7) suggests that biological neurons may use a subspace of the theoretically possible and much larger space of gene expression fitting their electrical phenotype. A putative explanation for this fact is that co-varying solutions might be much easier to generate (for instance through transcription regulatory mechanisms targeting ion channels belonging to a same co-expression modules) and might significantly decrease the expense of energy necessary to achieve a target phenotype.

Another potential source of redundancy dominance comes from our observation that models derived through repeated optimization around fixed biological targets ^26^ exhibit increased redundancy despite matching the same electrophysiological features. A similar redundancy-dominated structure was evident in our analysis of ion channel mRNA from excitatory neurons ^23^. In both cases, oversampling within local basins of the parameter or gene expression space leads to clustered solutions, further shaping the redundancy–synergy balance.

Our findings also indicate that excitability can emerge from multiple distinct configurations of underlying parameters, consistent with the principle of degeneracy. In this light, synergy observed among ion channels does not necessarily imply fragility, but may reflect this degeneracy: high-dimensional, flexible solution spaces that support stable excitability through diverse compensatory strategies. Such parameter interdependencies have been extensively described in modeling studies ^36,37,38,39,40,41^and are thought to underlie homeostatic regulation in real neurons, allowing stable function despite genetic variability, developmental changes, or perturbations ^42^. Thus, degeneracy, as revealed by synergistic parameter combinations, may be a key design principle of robust neural excitability.

While our study demonstrates the significant predictive power of ion channel genes in distinguishing interneuron types, it is important to acknowledge that a comprehensive understanding of neuronal diversity requires consideration of synaptic genes and other groups of genes. Recent research has highlighted the strong predictive power of synaptic and GPCR genes in defining neuronal subtypes and their electrical properties ^3^.

This research contributes to the growing body of evidence linking mRNA expression to observable neuronal phenotypes ^1,15,7^. By integrating computational modelling, statistical analysis and transcriptomic data, we provide a framework for understanding the complex relationships between gene expression and the electrophysiological properties that define neuronal function. This approach holds significant potential for future studies investigating the molecular basis of neuronal diversity and the pathogenesis of neurological disorders arising from disrupted neuronal signalling.

## Methods

### Detailed biophysical models

We constructed conductance-based models of mouse cortical pyramidal neurons and interneurons using morphologically reconstructed cells and a set of predefined ion channel mechanisms in NEURON environment^43^, as previously described and available online ^19^. Electrophysiological features were extracted from Patch Clamp recordings across various stimulation protocols using the eFEL library ^19,44^. Each model was optimized to match the canonical electrophysiological profile—defined as the mean feature vector of a given cell type—using a z-score–based cost function and a Markov Chain Monte Carlo (MCMC) procedure ^20^. Simulations were performed in NEURON, with ion channel distributions assigned by compartment and e-type. The resulting model populations included over 63,408 cAC, 86,361 bNAC, and 110,128 L5PC models that satisfied optimization thresholds (3 or 5 standard deviations). Full methodological details, including protocols, features, and channel distributions, are provided in the SI Appendix.

### Information theory measures and computations

We quantified high-order dependencies using the O-information (O-info) and the Redundancy–Synergy Index (RSI). O-info captures net redundancy or synergy among a set of variables **X** = {*X*_1_*, …, X_N_*}, based on their joint and marginal entropies ^21^. RSI measures the net interaction among predictors *S* relative to a target variable *Y*, defined as the difference between joint and individual mutual information terms ^22^. Both metrics were computed under a multivariate Gaussian assumption, enabling closed-form solutions based on the covariance matrix. Full definitions and derivations are provided in the SI Appendix.

### Gaussian formula for *O*-info

For a multivariate Gaussian system with covariance matrix Σ, the *O*-info is approximated as:

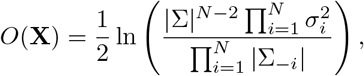

where:

- *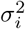* is the variance of the *i*-th variable (diagonal of S),
- |S| is the determinant of the covariance matrix.

### Gaussian formula for RSI

For RSI under Gaussian assumptions, the mutual information terms *I*(*S*; *Y*) and *I*(*X_i_*; *Y*) are calculated directly from the covariance matrices:

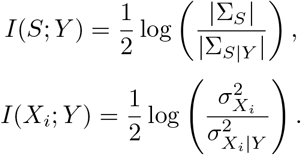

Substituting into the RSI formula:

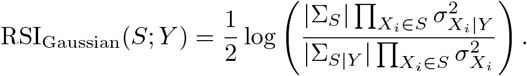

### Detection of Stabilization Points

To identify stabilization points in O-info or RSI curves, we applied a derivative-based algorithm. For each tuple order, we computed first and second discrete derivatives of the O-info/RSI series and defined the stabilization point as the index with the steepest concavity (minimum second derivative). This marks the transition from rapid decay to plateau. Full derivative expressions and rationale are provided in the SI Appendix.

### Clustering and analysis of the single cell mRNA from Patch-seq data

We analyzed single-cell RNA-seq data using Scanpy ^45^, applying standard preprocessing, log-transformation, and gene-wise standardization. Gene subsets included ion channel genes, differentially expressed genes (DEGs), and 1000 random sets of matched size. PCA was used for dimensionality reduction, followed by kNN graph construction and Leiden clustering. Cluster quality was assessed using Adjusted Rand Index (ARI) against known cell types. Full details on gene selection, normalization, and evaluation appear in the SI Appendix.

### Selection of the High-Covariance Model Subset

To assess the role of structured parameter co-variation, we selected a high-covariance subset from the cAC model ensemble (cost < 3) using a PCA-based method. Models were projected onto the top five principal components (explaining 70% of variance), reconstructed, and ranked by residual L2 error. The 1000 models with lowest reconstruction error were retained. The complete selection procedure is described in the SI Appendix.

### Mutual Information Computation and Significance Testing

We computed pairwise mutual information (MI) between parameters and electrophysiological features using histogram-based discretization. For each pair, significance was assessed by shuffling one variable (1000 permutations) and comparing to a null distribution. P-values were corrected for multiple comparisons using Benjamini–Hochberg FDR (*alpha* = 0.05). Only MI values passing FDR and a minimum effect size threshold (MI > 0.1) were considered significant. Details are provided in the SI Appendix.

### Bootstrap Estimation of high order measurements

To estimate confidence intervals for O-info and RSI metrics, we performed 50 bootstrap repetitions per condition. Samples of 1000 models (for MCMC data) or 60% of cells (for transcriptomic data) were drawn without replacement. Metrics were computed across tuple orders to capture variability. Datasets with fewer than 100 samples or strongly imbalanced clusters were excluded due to resampling artifacts. Full rationale and bootstrapping parameters are given in the SI Appendix.

## Data and Code Availability

All transcriptomic datasets are publicly available. Single-cell RNA-seq data from Tasic et al. (2018) and Patchseq data from Gouwens et al. (2020) can be accessed via the Allen Brain Map portal: https://portal.brain-map.org. Conductance-based models from Nandi et al. (2022) are available at: https://github.com/AllenInstitute/All-active-Manuscript. Code for computing O-information and RSI is provided at: https://github.com/BlueBrain/emodel-generalisation.

## Author Contributions

conceptualization, M.R., A.A., M.Z., JM.G.; methodology, M.R., A.A., A.M.; formal analysis, M.R., A.A.; investigation, M.R.; writing – original draft, M.R.; writing – review & editing, M.R., A.A., M.Z., A.M., JM.G., W.V.G; funding acquisition, H.M.. No competing interest.

## Acknowledgments

This study was supported by funding to the Blue Brain Project, a research center of the École polytechnique fédérale de Lausanne (EPFL), from the Swiss government’s ETH Board of the Swiss Federal Institutes of Technology. Authors would like to thank Dr. Fernando Rosas from the University of Sussex for the initial discussions and suggestions regarding IT tools.

## SI Methods

### Detailed biophysical models

Detailed biophysical model was constructed based on the patch clamp electrophysiological recordings of L5PC, cAC and bNAC cells as previously described and available online [1]. In the experimental dataset [1, 2], each recorded cell was probed with a series of stimulation protocols: (1) IDrest, consisting of depolarizing steps with a sampling frequency of 10 kHz and duration of 2 seconds; (2) IDthresh, also with depolarizing steps at a 10 kHz sampling frequency and 2-second duration; (3) APWaveform, with depolarizing steps at a higher sampling frequency of 50 kHz for 50 ms; (4) IV, a sequence of current steps ranging from hyperpolarization to depolarization, sampled at 10 kHz over 3 seconds. A holding current was applied to maintain the membrane potential at −70 mV (prior to a liquid junction potential correction of 14 mV). In total, the features extracted from the recordings include voltage measurements, such as voltage after stimulus, baseline voltage, maximum voltage relative to baseline, and voltage deflections during stimulus onset and o”set. Action potential (AP) properties, including AP amplitude, specific amplitudes (AP1, AP2, and last AP), AP half-width, and after-hyperpolarization (AHP) depth. Firing characteristics, such as mean firing frequency, total spike count, burst number, and ISI coe#cient of variation. Timing-related features, such as time to first and last spikes and the inverse timing of successive ISIs, further describe the temporal aspects of neuronal firing. Additionally, resistance and current measurements (input resistance, holding current, and threshold current) and the decay time constant following stimulus were assessed to characterize the cell’s intrinsic properties. The full description of the feature can be found at https://efel.readthedocs.io/en/latest/eFeatures.html.. These features were extracted for step protocols, the intensity of the step protocols reflects the fraction of the rheobase current (current necessary to elicit a single spike). The full list of protocols and features is can be found in SI.Table 1.

**Table 1:** Relevant protocols and e-features for bNAC, cAC, and cADPYR e-types in a single column format.

|  |
| --- |
| **E-type: bNAC, cNAC** |
| **Protocol: IDThresh/IDrest 150, 200, 250, 300 %**<br>E-features: voltage_base, voltage_after_stim, AP_amplitude, APlast_amp, AHP_depth, inv_time_to_first_spike, time_to_last_spike, inv_first_ISI, inv_second_ISI, inv_third_ISI, inv_fourth_ISI, inv_fifth_ISI, mean_frequency |
| **Protocol: APWaveform 360 %**<br>E-features: AP_amplitude, AP1_amp, AP_duration_half_width, AHP_depth |
| **Protocol: IV -100 %**<br>E-features: voltage_deflection, voltage_deflection_begin |
| **Protocol: IV -20 % (Rin)**<br>E-features: ohmic_input_resistance_vb_ssse, voltage_base |
| **Protocol: IV 0 % (RMP)**<br>E-features: voltage_base, Spikecount |
| **Protocol: RinHoldCurrent**<br>E-features: bpo_holding_current |
| **Protocol: Threshold**<br>E-features: bpo_threshold_current |
| **E-type: cAC** |
| **Protocol: IDThresh/IDrest 140, 200, 250, 300 %**<br>E-features: voltage_base, voltage_after_stim, AP_amplitude, APlast_amp, AHP_depth, inv_time_to_first_spike, time_to_last_spike, inv_first_ISI, inv_second_ISI, inv_third_ISI, inv_fourth_ISI, inv_fifth_ISI, mean_frequency |
| **Protocol: APWaveform 360 %**<br>E-features: AP_amplitude, AP1_amp, AP_duration_half_width, AHP_depth |
| **Protocol: IV -100 %**<br>E-features: voltage_deflection, voltage_deflection_begin |
| **Protocol: IV -20 % (Rin)**<br>E-features: ohmic_input_resistance_vb_ssse, voltage_base |
| **Protocol: IV 0 % (RMP)**<br>E-features: voltage_base, Spikecount |
| **Protocol: RinHoldCurrent**<br>E-features: bpo_holding_current |
| **Protocol: Threshold**<br>E-features: bpo_threshold_current |
| **E-type: cADPYR** |
| **Protocol: APWaveform 320 %**<br>E-features: AP_amplitude, AP1_amp, AP2_amp, AP_duration_half_width, AHP_depth |
| **Protocol: IV -100 %**<br>E-features: voltage_deflection, voltage_deflection_begin |
| **Protocol: IDrest & IDthresh 150, 200, 280 %**<br>E-features: voltage_base, voltage_after_stim, AP_amplitude, APlast_amp, AHP_depth, inv_time_to_first_spike, time_to_last_spike, inv_first_ISI, inv_second_ISI, inv_third_ISI, inv_fourth_ISI, inv_fifth_ISI, mean_frequency |
| **Protocol: SpikeRec 600 %**<br>E-features: decay_time_constant_after_stim, voltage_after_stim, Spikecount |
| **Protocol: IV -20 % (Rin)**<br>E-features: ohmic_input_resistance_vb_ssse, voltage_base |
| **Protocol: IV 0 % (RMP)**<br>E-features: voltage_base, Spikecount |
| **Protocol: RinHoldCurrent**<br>E-features: bpo_holding_current |
| **Protocol: Threshold**<br>E-features: bpo_threshold_current |

The neuronal models were implemented in NEURON simulator [3]. We used a set of ionic mechanisms described previously, the table with mechanisms specific for each electrical type can be found in SI.Table 2.

**Table 2:** Active parameters and their compartmental placement for cAC, bNAC, and cADPYR e-types.

|  |
| --- |
| **E-type: cADPYR** |
| **Soma**<br>CaDynamics, HVA Ca, LVA Ca, Kv3.1, Ca-activated K, Persistent K, Transient K, Transient Na<br>**AIS**<br>CaDynamics, HVA Ca, LVA Ca, Kv3.1, Ca-activated K, Persistent K, Transient K, Transient Na, Persistent Na<br>**Dendrites**<br>CaDynamics, HVA Ca, LVA Ca, Kv3.1 (Apical), Transient Na (Apical; Decaying), HCN (Apical and Basal; Exponential increasing towards terminals) |
| **E-types: bNAC, cAC** |
| **Soma**<br>CaDynamics, HVA Ca, LVA Ca, Ca-activated K, Transient Na, Kv3.1, Persistent K, Transient K, HCN<br>**AIS**<br>CaDynamics, HVA Ca, LVA Ca, Ca-activated K, Transient Na, Kv3.1, Persistent K, Transient K<br>**Dendrites**<br>CaDynamics, HVA Ca, LVA Ca, Ca-activated K, HCN (Exponential increasing towards terminals) |

Morphologies were manually reconstructed. The axons and their branches were replaced by a synthetic axon section consisting of an AIS (60 *u*m) followed by a myelinated axon segment of 1,000 *u*m. To define the optimization target for each cell type, we computed a feature vector representing the mean electrophysiological properties across multiple recorded neurons. As such, each biophysical model in our study is optimized to match this canonical feature set, rather than reproducing variability across biological replicates. To optimize these models, we employed an MCMC procedure as described in [4], with the cost computed as:

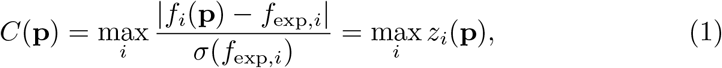

where *z_i_* are the absolute z-scores for each feature *i* and simulated feature values are represented as a vector **f** = (*f*_0_*,..., f_k_*), and the parameters are denoted as **p** = (*p*_0_*,..., p_n_*).

From the resulting parameter sets, we selected models with an optimization cost within 3 standard deviations for cAC and bNAC, yielding 63,408 cAC models and 86,361 bNAC models. For L5PC, we selected models with an optimization cost within 5 standard deviations, resulting in 110,128 L5PC models.

### Detection of Stabilization Points

To identify the stabilization point in the O-info (or RSI) value series, we employed a derivative-based algorithm. This approach utilizes the first and second derivatives of the RSI curve to detect the point where the data stabilizes, as follows:

- **Data Extraction:** For each tuple, the maximum RSI value is extracted, forming a series of values *y* = {*y*_1_*, y*_2_*,..., y_n_*} corresponding to tuple indices *x* = {1, 2*,..., n*}.
- **Derivative Calculation:** The first derivative *y*(*x*) is computed as the rate of change between consecutive RSI values:

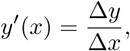

where Δ*y* = *y*_*i*+1_*y_i_* and Δ*x* = *x*_*i*+1_ *x*_*i*_. Subsequently, the second derivative *y*(*x*) is calculated to capture the rate of change in the slope:

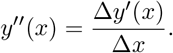
- **Stabilization Point Detection:** The stabilization point is identified as the index where the second derivative *y*(*x*) attains its most negative value, corresponding to the steepest concave point in the RSI curve.

Mathematically:

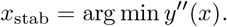

This point marks the transition from steep decline to stabilization in the curve.

### Clustering and analysis of the single cell mRNA from Patchseq data

Data preprocessing and clustering were performed on single-cell RNA sequencing data containing expression profiles for thousands of genes across cells [5]. Ground truth cell type labels were used to evaluate clustering performance. The analyses and clustering were implemented using Python libraries, scanpy [6] was used for bio-informatics analysis (preprocessing and clustering workflows).

Gene subsets were selected as follows:

- Ion Channel Subset: A predefined list of ion channel genes was used to extract a subset of the expression matrix.
- Di”erentially Expressed Gene (DEG) Subset: DEGs were identified using statistical tests (e.g., t-tests) to compare expression levels across conditions.
- Random Subsets: Control subsets were generated by selecting 1,000 random gene sets, each matching the size of the ion channel subset.

Normalization was applied to ensure consistency in expression data:

- Log-transformation (*log*_2_(*counts* + 1)) was used to normalize skewed distributions.
- Standardization was performed to scale each gene’s expression to have zero mean and unit variance.

Dimensionality reduction was conducted using Principal Component Analysis (PCA) to reduce data dimensionality while retaining features explaining most of the variance (e.g., 90%).

Clustering was performed using the Leiden algorithm, which is optimized for detecting communities in graphs derived from high-dimensional data. Key steps included:

- Graph Construction: A k-nearest neighbor (kNN) graph was constructed from the principal components, with edges indicating cell similarities.
- Community Detection: The Leiden algorithm partitioned cells into clusters, with the number of clusters determined adaptively from the graph structure.
- Evaluation: Clustering results were compared to ground truth cell type labels using the Adjusted Rand Index (ARI). ARI measures label similarity, correcting for chance, with a score of 1 indicating perfect agreement.

Clustering was used to assess the specificity of ion channel, randomly selected and DEG subsets. The same preprocessing and clustering steps were repeated for 1,000 random gene subsets. ARI scores were computed for each run and compared to the ground truth labels. The resulting distribution of ARI scores served as a baseline for comparison.

### Selection of the High-Covariance Model Subset

To assess how structured co-variation among model parameters influences high-order interactions, we identified a subset of models characterized by elevated inter-parameter dependencies using a principal component analysis (PCA)-based selection strategy.

Starting from the full ensemble of valid conductance-based models for cAC interneurons (i.e., models with cost ¡ 3), we performed PCA on the matrix of normalized parameters (20 per model). We retained the top N = 5 principal components, which captured 70% of variance across the parameter space. We then reconstructed each model from its projection onto these top components and quantified the residual error between the original and reconstructed parameter vectors using the L2 norm. We ranked all models by their reconstruction error and selected the top 1000 models with the lowest residuals as our high-covariance subset.

### Mutual Information Computation and Significance Testing

To quantify the statistical dependency between model parameters and extracted features, we computed pairwise mutual information (MI) using a non-parametric estimator. For each parameter-feature pair, we estimated MI based on histogram-based discretization. To assess statistical significance, we applied a bootstrap procedure for each MI value. Specifically, we generated a null distribution by independently shu!ing one variable in the pair (1000 permutations), and recomputing the MI to obtain an empirical p-value. These p-values were collected into a matrix and subsequently corrected for multiple comparisons using the Benjamini–Hochberg procedure (FDR control at *ε* = 0.05). After correction, only MI values that passed the FDR threshold and exceeded a minimal e”ect size threshold of 0.1 were considered significant and visualized (e.g., in the network graph and circos plots).

### Bootstrap Estimation of high order measurements

To assess the robustness and variability of higher-order information metrics (O-information and Redundancy-Synergy Index, RSI), we implemented a bootstrap resampling procedure. Specifically, we drew 50 independent bootstrap samples, each consisting of 1000 (unless stated di”erently) for the MCMC generated models or 60% of the total sample number for the genetics data selected without replacement from the original model population for each cell type. For each resampled dataset, we computed O-information or RSI across tuple orders. This allowed us to estimate the variability of maximum and minimum values per order and to derive 95% confidence intervals from the bootstrap distributions (based on the 2.5th and 97.5th percentiles). In the figures, we plot all individual bootstrap curves along with shaded confidence envelopes, providing a quantitative assessment of the stability of synergy- or redundancy-dominated regimes. This procedure was applied uniformly across all datasets analyzed in the manuscript, including both biophysical model populations and transcriptomic data, to enable direct comparison of the robustness of inferred information-theoretic structures. However, we did not apply bootstrapping to datasets with fewer than 100 samples, or to data that includes heterogeneous clusters with small sample sizes (e.g., 10–30 samples), due to the risk of distortion from resampling limited or unevenly distributed data. In such cases, bootstrapping can over-or underrepresent minor clusters, disrupt underlying population structure, and yield unreliable estimates of high-order interactions. Given the sensitivity of O-info to subtle correlation patterns, we instead relied on direct estimation using the full available data and reported values without confidence intervals.

---

**Figure S1:**
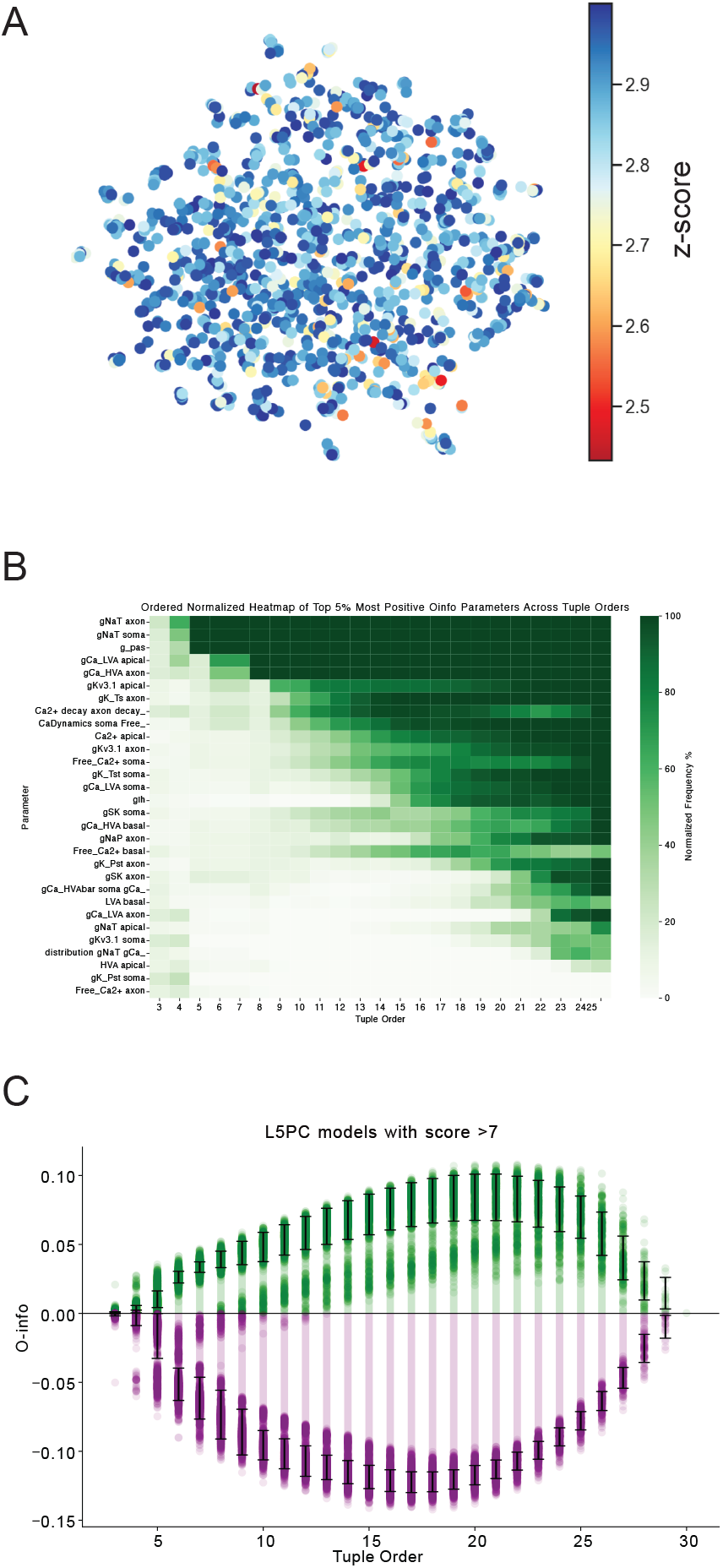
**A.**UMAP representation of model parameters. The colors indicate the fitness of the model, quantified as the z-score for the corresponding parameter set. B. Normalized frequency map highlighting parameters that frequently occur in high-redundancy tuples. C. O-information plot showing the relationship between tuple number (number of parameters) and O-information values for models L5PC models that have score value larger than 7. Each dot represents the O-information value for a specific tuple. The bars and whiskers illustrate the mean and standard deviation of O-information values at each tuple order. The color encodes positive (green) and negative (purple) O-info values.

**Figure S2:**
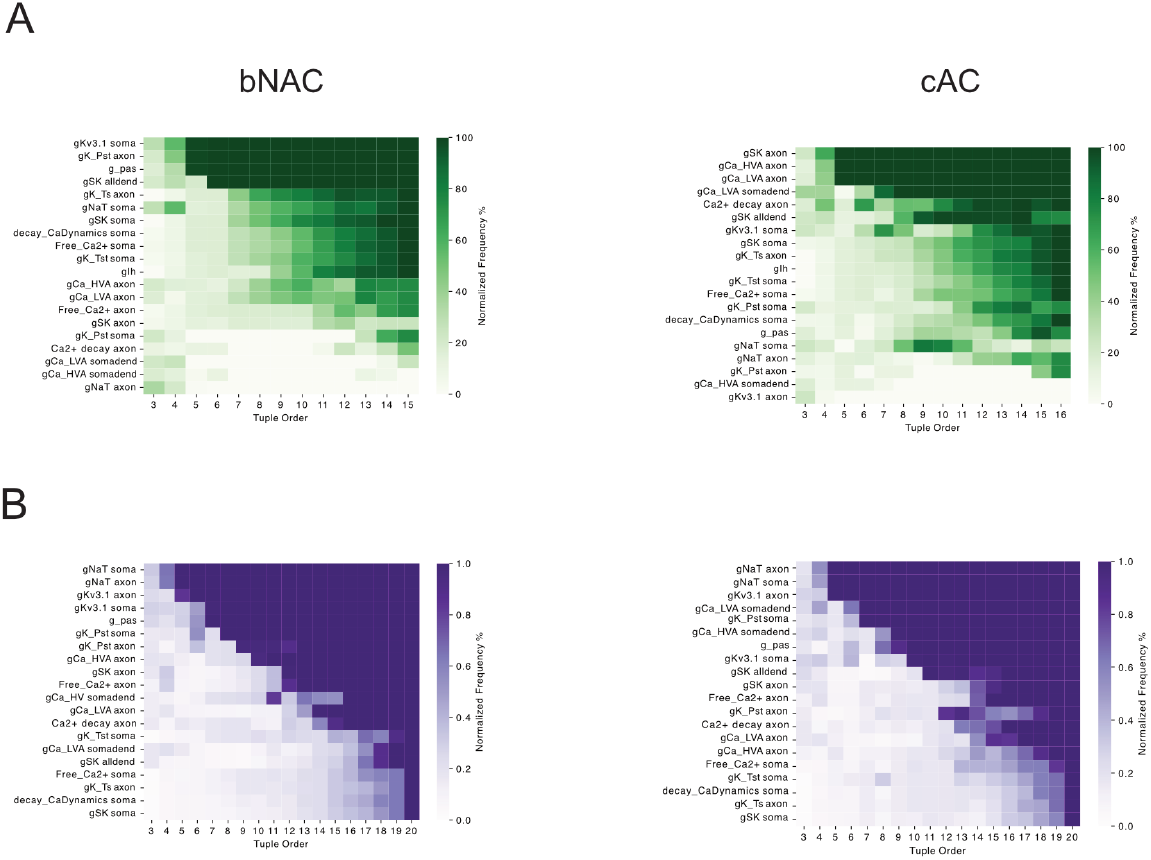
**A.**Normalized frequency map showing parameters that frequently appear in high-redundancy tuples based on Oinfo calculations for cAC and bNAC neuronal models. B. Normalized frequency map for parameters in highsynergy tuples, similar to panel.

**Figure S3:**
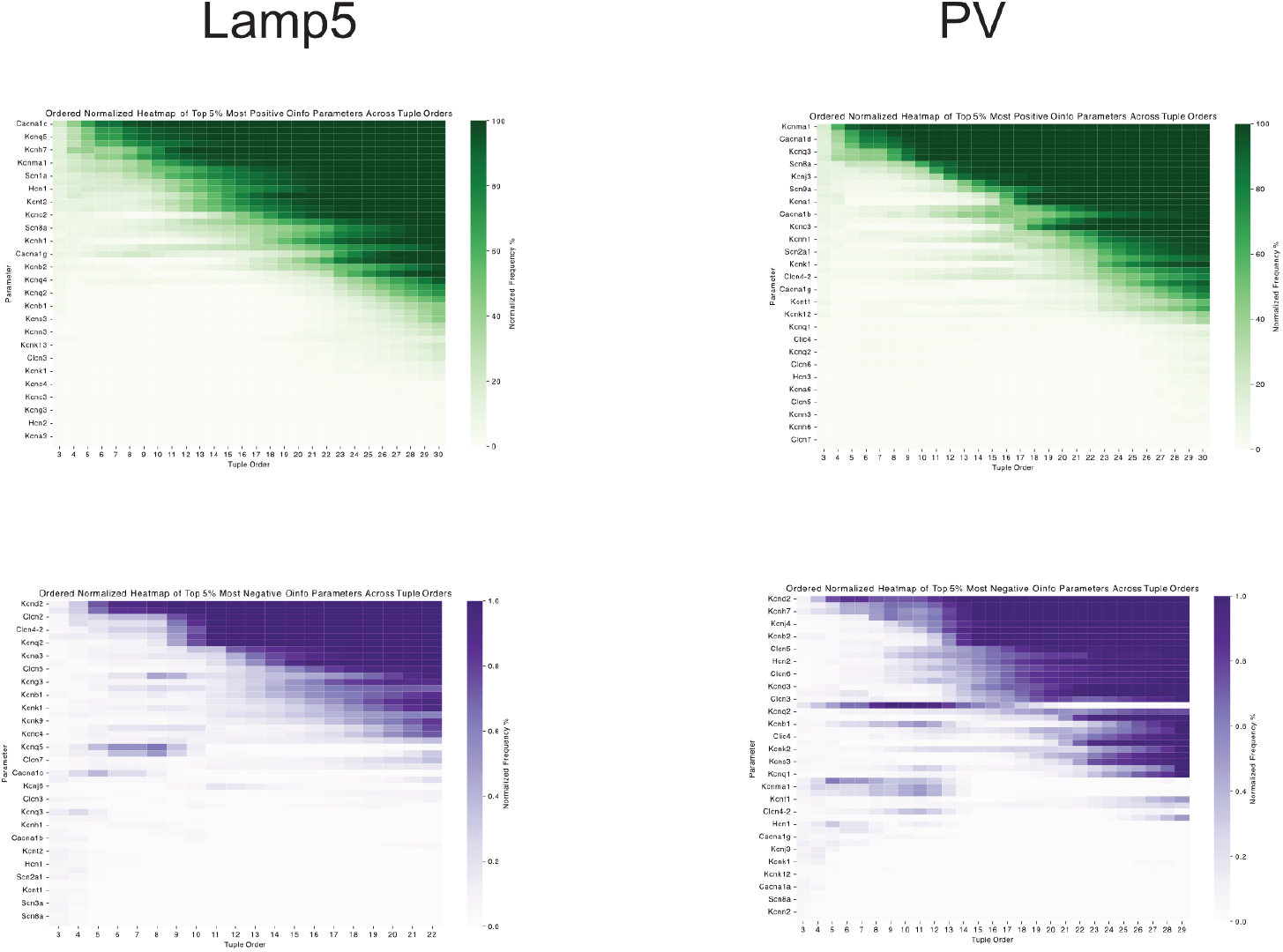
Normalized frequency maps of parameters frequently appearing in high-redundancy (top) and high-synergy (bottom) tuples for Lamp5 and Pvalb neurons.

**Figure S4:**
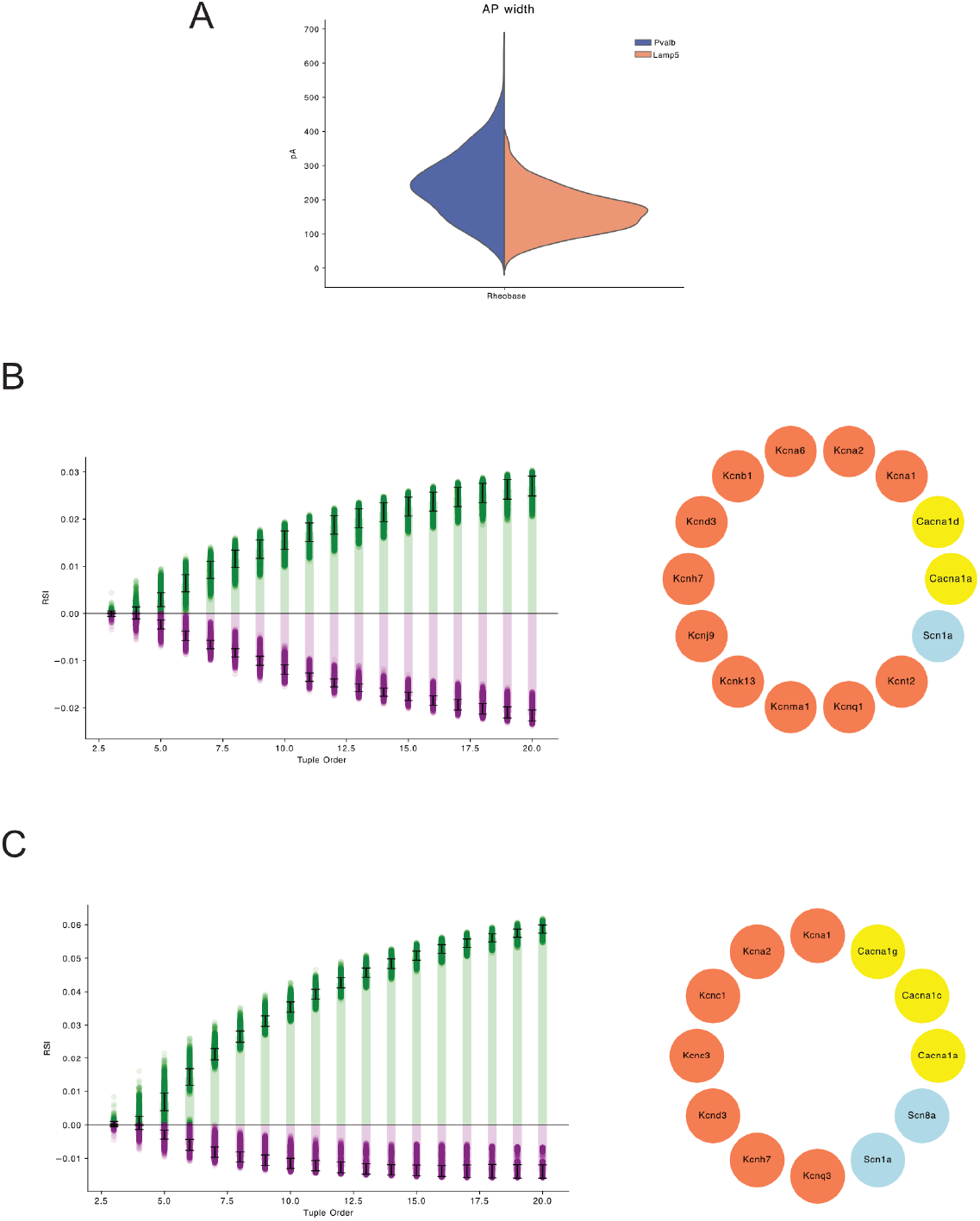
**A.**Distribution of mean frequencies (Hz) for cells from Lamp5 and Pvalb neuronal types. B. RSI values plotted against tuple numbers (right), along with the set of genes that yield the highest RSI value (left) for Lamp5 neurons. C. Similar RSI values and gene sets as in panel B, but for Pvalb neurons.

**Figure S5:**
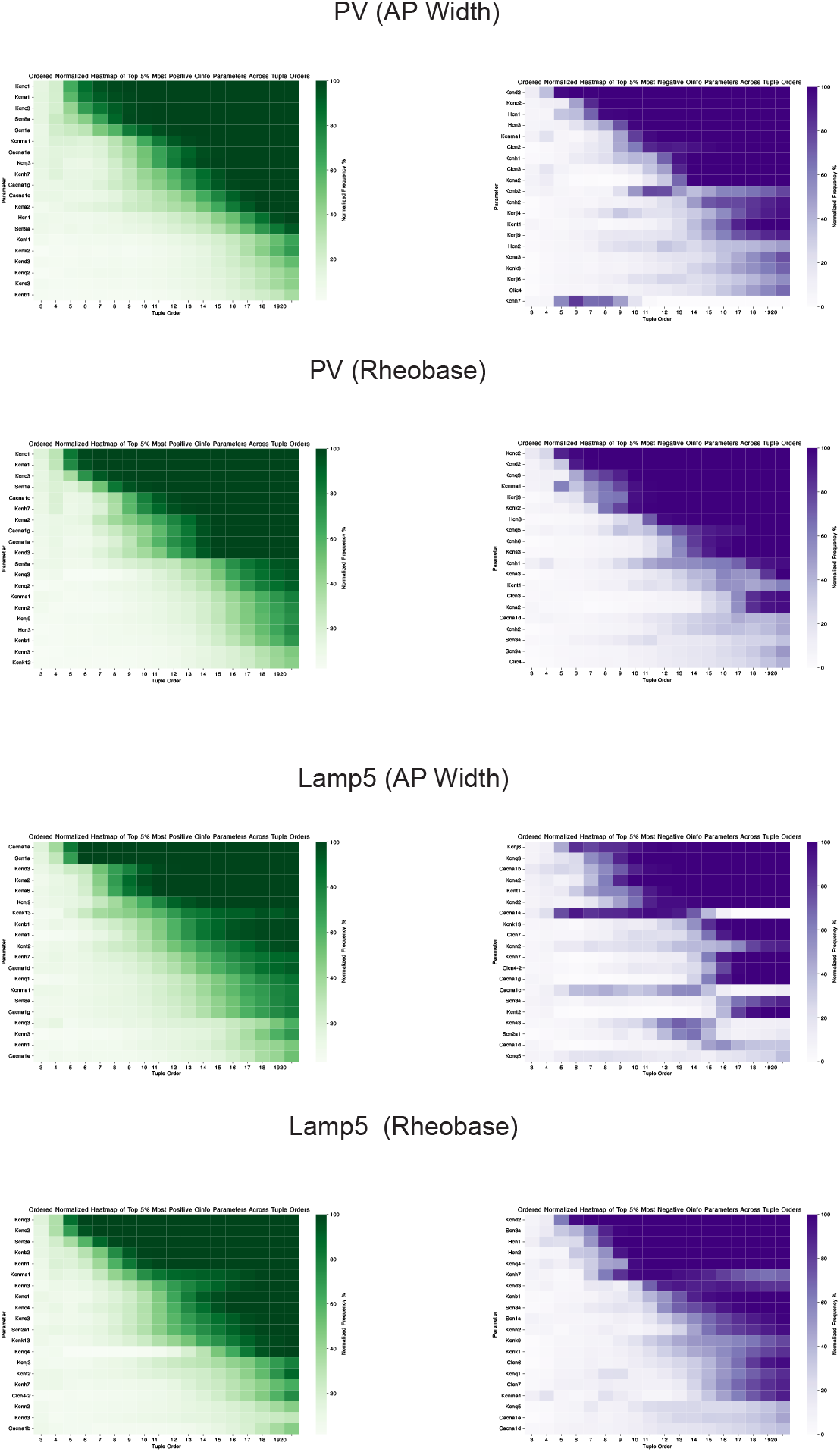
Normalized frequency maps of parameters frequently appearing in high-redundancy (left) and high-synergy (right) tuples. Data correspond to gene expression, action potential width (AP width), and mean frequency features for Lamp5 and Pvalb neurons.

**Figure S6:**
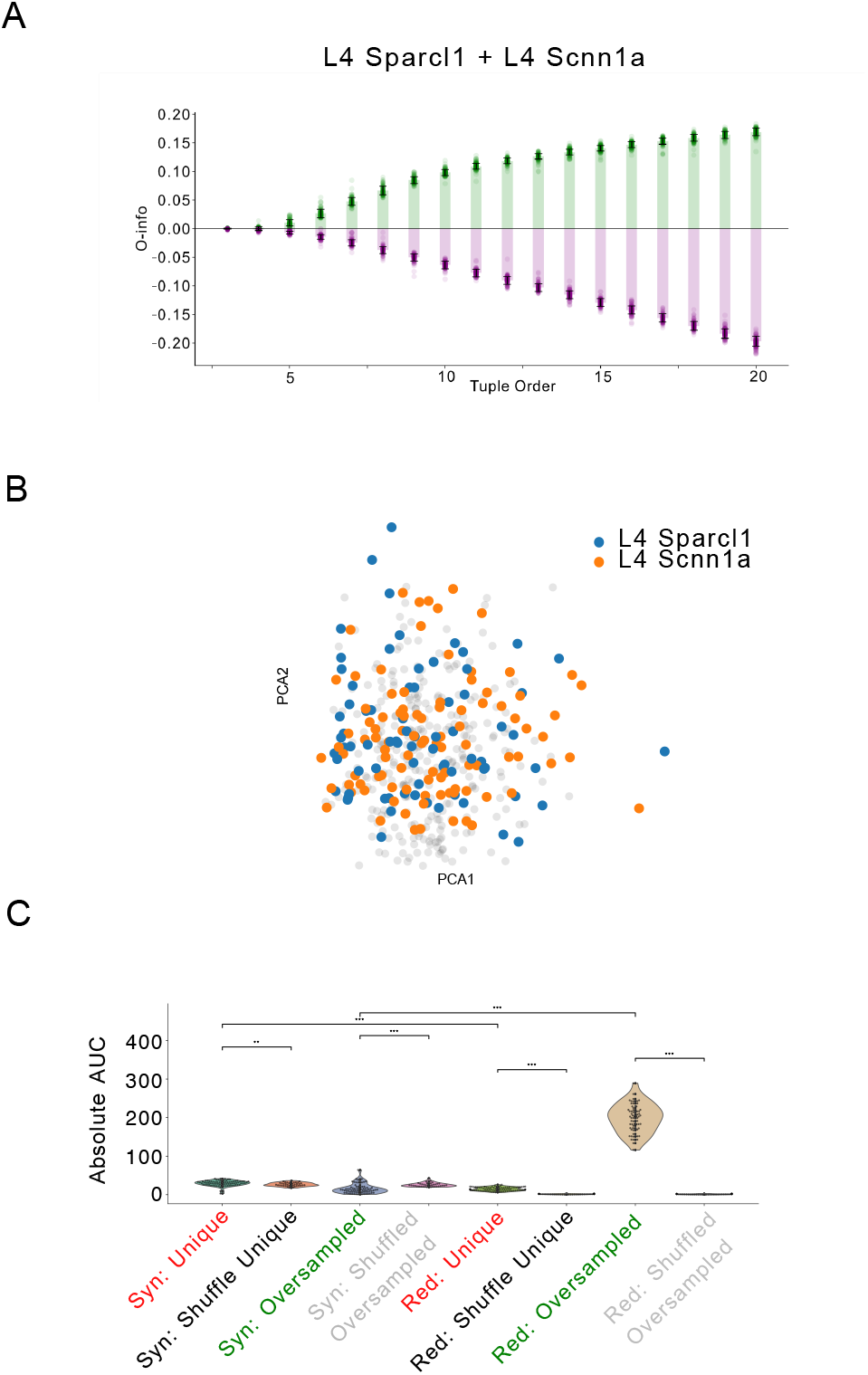
**A.** O-info decomposition restricted to models from the L4 Sparcl1+ and L4 Scnn1a+ excitatory neuron subtypes. Green and purple curves represent the mean ± SD of positive and negative O-information values, respectively, as a function of tuple order. **B.** Projection of the parameter sets corresponding to L4 Sparcl1+ and L4 Scnn1a+ gene expression onto the first two principal components. Colors indicate subtype identity. **C.** Distribution of absolute O-info AUC values for both synergy (minima) and redundancy (maxima) across the Unique, Oversampled, and Shu!ed groups. Violin plots display group-wise variability, with statistical comparisons performed using Mann-Whitney U tests (p < 0.001).

